# Novel GαGTP Sensors Reveal Endogenous and Subcellular G Protein Signaling Dynamics

**DOI:** 10.64898/2026.01.29.702668

**Authors:** Dhanushan Wijayaratna, Senuri Piyawardana, Ajith Karunarathne

**Affiliations:** Department of Chemistry, Saint Louis University, Saint Louis, MO 63103; Institute for Drug and Biotherapeutic Innovation, Saint Louis University, Saint Louis, MO 63103; Division of Gastroenterology, Hepatology and Nutrition, Boston Children’s Hospital; Harvard Medical School, Boston, MA 02115

**Keywords:** G proteins, GPCR, Sensors, Subcellular, Membrane-targeting, Signaling Adaptation

## Abstract

G protein-coupled receptors (GPCRs) perceive spatially and temporally diverse stimuli and activate G protein heterotrimers comprising α, β, and γ subunits, which broadcast signals through a broad range of effectors at various subcellular compartments. Therefore, understanding endogenous G protein activity dynamics at the subcellular level, thereby recapitulating in vivo signaling paradigms, will facilitate the identification of pathological signaling pathways. However, the lack of sensors for endogenous G proteins has been an obstacle. Here, we demonstrate the engineering of sensors to probe endogenous GαiGTP and GαqGTP. Compared to examining overexpressed and fluorescently tagged Gα, our sensors capture the magnitude and kinetics of endogenous GαGTP dynamics, including their generation, equilibrium signaling, and hydrolysis, with native fidelity. Using the translocation-based GαiGTP sensor, we show that heterotrimer dissociation upon Gi-GPCR activation is Gγ-subtype dependent. Confirming our previous findings, the GαqGTP sensor showed that Gαq expression is low and tightly regulated in most cells. Using optogenetic tools, we demonstrate that our sensors detect GαGTP generation and hydrolysis during asymmetric GPCR-G protein activation, a capability that will be particularly useful in morphologically diverse cells such as neurons. Therefore, our engineered novel GαGTP sensors can be highly beneficial in decoding subcellularly resolved endogenous G protein signaling dynamics.

## Introduction

GPCRs constitute the largest class of cell-surface receptors in humans,^1-3^ activated by hormones, peptides, neurotransmitters, lipids, and light^4-6^ to initiate intracellular signaling pathways through G protein activation, which in turn induces a wide array of cellular and physiological responses. G protein heterotrimers comprise α and βγ subunits, transmitting signals from GPCRs to the cell interior.^5,7,8^ G protein pathways can be grouped based on the type of Gα subunit in the heterotrimer. Gα proteins are grouped into four families: Gα_s,_ Gα_q,_ Gα_i,_ and Gα_12/13_ ^9^ In the inactive GPCR form, Gα is bound to GDP, and upon GPCR activation, conformational changes in Gα allow for spontaneous GDP to GTP exchange in Gα, which induces Gβγ dissociation from GαGTP.^10^ The resultant GαGTP and free Gβγ interact with effector proteins to initiate downstream cellular signaling.^11^ The extensive involvement in cell signaling, physiology, and presence on the cell surface make GPCRs the major drug target.^12^ Due to the pharmacological significance, sensors/ probes for GPCR-G protein signaling are currently a major focus in the field.

Mini-G proteins are widely used for probing active-state GPCRs.^13^ Currently available sensors/ assays for G proteins include using FRET/ BRET-based fluorescently-tagged Gα and Gβγ dissociation,^14-17^ translocation-based fluorescently-tagged Gα and Gβγ,^16,18-20^ indirect sensors detecting downstream activity (sensors for cAMP, Ca^2+^, PIP2 hydrolysis, etc),^21-24^ and more recently developed TRUPATH assay,^25^ BERKY,^26^ and ONE-GO bio-sensors.^27^ Most of these above-mentioned assays (FRET/ BRET assays, translocation-based assays, and TRUPATH assay) require the overexpression of fluorescently-tagged G proteins.^28,29^ We have previously shown that genetic modification of cells by overexpression of G proteins can drastically change signaling extents and kinetics.^30,31^ On the other hand, BERKY and ONE-GO biosensors that can detect endogenous GαGTP represent a vast improvement over the previous sensors. However, since they are BRET sensors, they cannot provide dynamic subcellular information on endogenous G protein activation. Therefore, it is imperative to use sensors that can detect active endogenous G proteins at the subcellular level.

Gq and Gi/o pathways signal through distinct downstream effectors.^32^ Active Gαq (GαqGTP) activates phospholipase Cβ (PLCβ), which hydrolyzes phosphatidylinositol 4,5-bisphosphate (PIP2) to diacylglycerol (DAG) and inositol triphosphate (IP3).^33^ IP3 promotes Ca^2+^ release from endoplasmic reticulum (ER) calcium stores through IP3 receptor (IP3R) channels non-energetically.^34^ DAG activates protein kinase C (PKC).^33^ Cytosolic Ca^2+^ increase together with PKC activation regulates calcium-regulated kinases,^35^ mitogen-activated protein kinases (MAPKs),^36^ which mediate cellular functions ranging from platelet activation and aggregation,^37^ cell proliferation,^36^ and secretion.^38^ Due to the regulation of downstream signaling inducing cell proliferation, the Gq pathway is considered an oncogenic pathway; therefore, it is tightly regulated in cells. Like GαqGTP, activation of Gi-coupled GPCRs results in GαiGTP formation, inhibiting adenylyl cyclase and terminating cAMP production, thereby inhibiting cAMP-dependent protein kinase activity.^39,40^ GαiGTP also promotes protein tyrosine kinase C-Src activity,^41^ which is involved in cell migration, metabolism, and differentiation.^42^

Although the type of Gα has been considered the primary determinant of the overall pathway function, the significance of Gβγ as a vital regulator has become apparent over the last two decades.^20,29^ We have established that the Gγ subunit modulates the plasma membrane affinity of the Gβγ since Gγ provides the only plasma membrane anchor for the Gβγ dimer.^29,43^ We have also shown that the Gγ subtype-dependent plasma membrane affinities of Gβγ control the dynamics and the efficacy of several effector pathways, including phospholipase C (PLC),^44^ Phosphoinositide 3-kinases (PI3K), GPCR kinases (GRKs).^29,31,45^ Additionally, evidence suggests Gβγ regulates adenylyl cyclases,^46^ inwardly rectifying potassium (GIRK) channels,^47^ Ca^2+^ channels,^48^ and guanine nucleotide exchange factors (GEFs).^49^ However, to our knowledge, no studies have explored how or whether the Gγ subunit in the Gβγ dimer influences GPCR-G protein activation-deactivation dynamics, mainly due to the lack of tools.

Here, we show the engineering of two sensors with unique designs that translocate to cell membranes, where GαiGTP and GαqGTP are generated upon GPCR activation. Using live-cell imaging, subcellular optogenetics, and single-cell analysis, we demonstrate the utility of our two sensors for detecting endogenous Gαi and Gαq-GTP generation and dynamics at various subcellular locations upon the onset of GPCR activity. Our results show that Gq signaling is tightly regulated in cells and that the Gγ subunit of Gβγ is a primary regulator of Gi-GPCR-mediated signaling efficacy. Since Gγ subtypes exhibit cell and tissue-specific expression patterns,^43^ the findings of this study provide molecular insights into the evolution of the 12-member Gγ family and how the predominantly expressed Gγ subtypes can influence the signaling paradigms of a cell.

## Results and Discussion

### Fluorescently tagged GαiGTP and GαqGTP-interacting peptides fail to show detectable interactions upon receptor activation

Our objective was to design peptide-based sensors that are cytosolic before GPCR-G protein activation and translocate to the plasma membrane to detect endogenous GαGTP, generated upon corresponding GPCR activation. We therefore sought peptides, KB1753 (for Gαi) and GRK2-RH (for Gαq), that reported to bind Gαi-GTP and Gαq-GTP with high affinity, while showing little to no interaction with the GDP-bound form of Gα as potential starting points.^26,28^ To test the feasibility, we generated simple versions of the sensors: Venus-tagged versions of KB1753 (KB1753-Venus) and GRK2-RH (GRK2-RH-Venus), and expressed in HeLa cells together with the CFP-tagged α2-Adrenergic Receptor (α2AR-CFP), a Gi/o-coupled GPCR,^50^ or Gastrin-Releasing Peptide Receptor (GRPR), a Gq-coupled GPCR,^51^ respectively. As expected, KB1753-Venus was cytosolic (Fig. 1a, 0 s), given that almost all Gαi subunits in cells before GPCR activation are GDP-bound, either in the heterotrimer or as free GαiGDP localized to the plasma membrane.^52^ This is due to the GTPase activities of the Gα subunit itself, as well as the small GTPases that are acting as signaling regulators.^5^ Next, we activated α2ARs using 100 µM norepinephrine (NE), which induces the GDP-to-GTP exchange in Gαi. However, we did not observe a detectable recruitment of the sensor to the plasma membrane (Fig. 1a, 90 and 240 s, and plot). Similarly, GRK2-RH-Venus was also localized to the cytosol (Figure 1d, 0 s), suggesting that GRK2-RH did not show a detectable affinity towards GαqGDP. Since the endogenous Gαq expression in most cell lines, including HeLa cells, is significantly lower compared to other Gα types (Gαi/o, Gαs),^30,31^ in addition to GRK2-RH-Venus and GRPR, we also expressed Gαq-CFP in HeLa cells, and activated GRPR using 1 µM bombesin. Despite the Gαq expression, we did not observe a measurable recruitment of GRK2-RH-Venus to the plasma membrane (Fig. 1d, 90 and 240 s, and plot).

**Figure 1.**
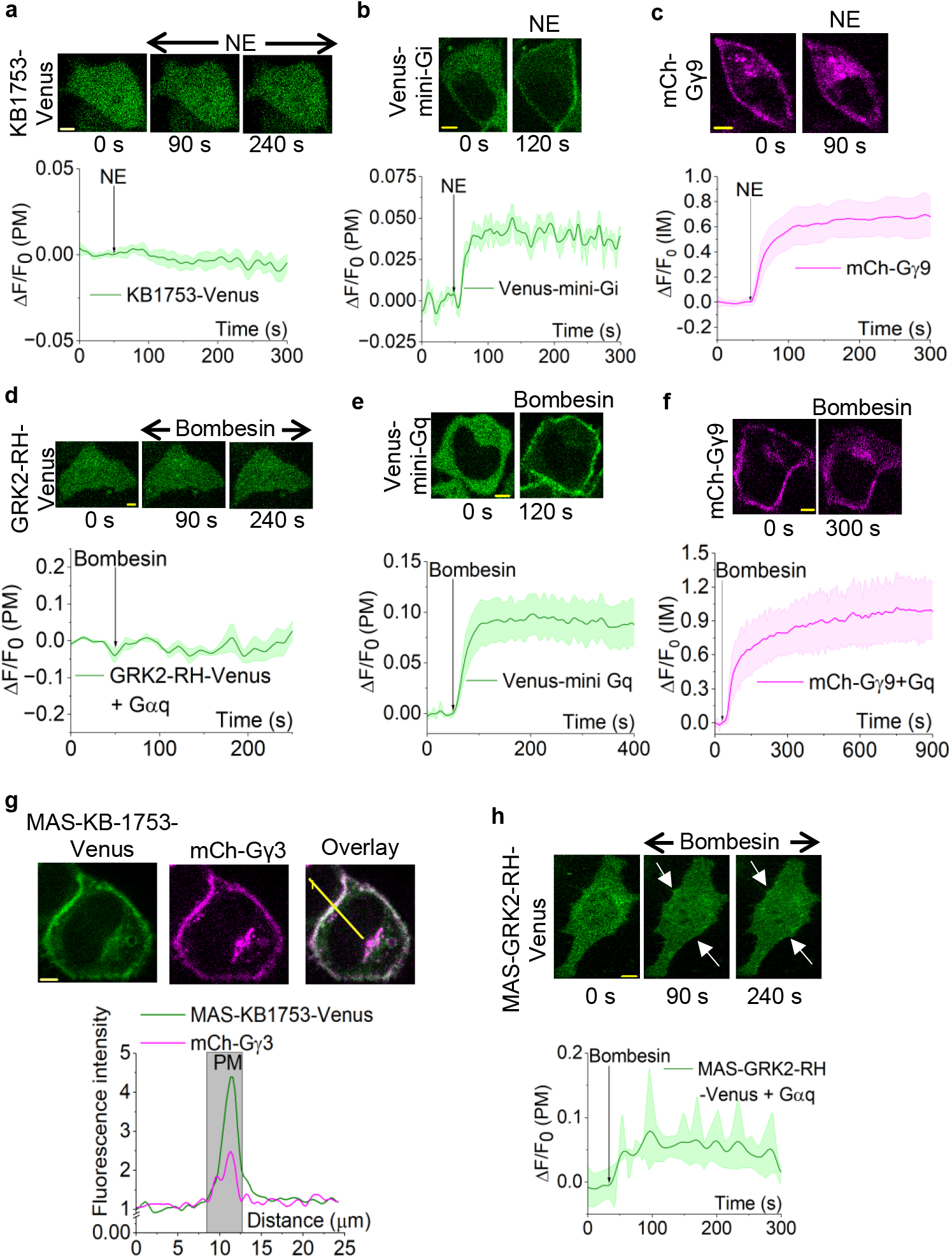
Fluorescently tagged KB1753 and GRK2-RH are inefficient probes for Gαi and GαqGTP, respectively. (a) Hela cells expressing α2AR-CFP and KB1753-Venus exhibit no recruitment to the plasma membrane upon α2AR activation with 100 μM norepinephrine. KB1753-Venus is localized to the cytosol before and after receptor activation. The corresponding plot shows the KB1753 dynamics on the plasma membrane (n = 9). (b) Upon α2AR activation in HeLa cells expressing α2AR-CFP and Venus-mini-Gi, mini-Gi is robustly recruited to the plasma membrane (120 s). Venus-mini-Gi plasma membrane dynamics are shown in the corresponding plot (n = 8). (c) HeLa cells expressing α2AR-CFP and mCh-Gγ9 showed robust Gγ9 translocation from the plasma membrane to internal membranes (90 s) upon α2AR activation with norepinephrine. The corresponding plot shows the mean mCherry fluorescence intensity profile in internal membranes (n = 10). (d) Hela cells expressing GRPR, GRK2-RH-Venus, and Gαq-CFP do not show any recruitment to the plasma membrane upon GRPR activation with 1 μM bombesin. The corresponding plot shows GRK2-RH-Venus dynamics on the plasma membrane (n = 10). (e) Robust mini-Gq recruitment to the plasma membrane was observed upon GRPR activation in HeLa cells expressing GRPR and Venus-mini-Gq (120 s). The corresponding plot shows Venus-mini-Gq dynamics on the plasma membrane (n = 10). (f) HeLa cells expressing GRPR, Gαq-CFP, and mCh-Gγ9 showed efficient Gγ9 translocation (300 s) upon GRPR activation. mCh-Gγ9 profile in internal membranes is given in the following plot (n = 13). (g) MAS-KB1753-Venus was predominantly localized to the plasma membrane in HeLa cells expressing α2AR-CFP and MAS-KB1753-Venus. Overlay between MAS-KB1753-Venus and plasma membrane-bound mCh-Gγ3 indicates that MAS-KB1753-Venus is near-completely localized to the plasma membrane. The line profile plot indicates the fluorescence intensity increase on the plasma membrane due to MAS-KB1753-Venus (green plot) and mCh-Gγ3 (magenta plot). (h) Hela cells expressing GRPR, MAS-GRK2(RH)-Venus, and Gαq-CFP exhibit cytosolic distribution of MAS-GRK2(RH)-Venus before GRPR activation (0 s). Upon GRPR activation with 1 μM bombesin, MAS-GRK2(RH)-Venus shows minor recruitment to the plasma membrane (white arrows). The corresponding plot shows MAS-GRK2(RH)-Venus dynamics on the plasma membrane (n = 9). Average curves were plotted using ‘n’ cells, n= number of cells. Error bars represent SEM (standard error of mean). The scale bar = 5 µm. NE: Norepinephrine; PM: Plasma membrane; IM: Internal membranes; ΔF: Fluorescence intensity change; F_0_: Baseline fluorescence

To ensure that the negligible probe recruitment is due to the inability of the probes to detect GαGTPs, and not due to defective receptor signaling or heterotrimer dissociation, we examined receptor activity and heterotrimer dissociation using mini-G sensor recruitment^13^ and Gβγ9 translocation,^43^ respectively. Mini-G sensor and Gβγ9 assay principles are depicted in the Fig. S1 diagram. Here, we expressed α2AR-CFP alongside Venus-mini-Gi and mCherry (mCh)-Gγ9 in HeLa cells. Upon activation of α2AR with 100 µM norepinephrine, we observed a robust recruitment of Venus-mini-Gi to the plasma membrane, indicating significant receptor activity (Fig. 1b) and robust mCh-Gγ9 translocation from the plasma membrane to internal membranes, indicating significant heterotrimer dissociation (Fig. 1c). In a separate experiment, we similarly examined GRPR activation using HeLa cells expressing Venus-mini-Gq, mCh-Gγ9, and Gαq-CFP. Upon GRPR activation with 1 µM bombesin, we also observed a strong recruitment of mini-Gq to the plasma membrane (Fig. 1e) and a robust Gγ9 translocation to internal membranes (Fig. 1f), indicating effective receptor activity and heterotrimer dissociation, respectively. This data indicated that the KB1753 and GRK2-RH peptides do not have sufficient GαGTP binding when expressed in the cytosol as simple sensors.

### Engineering N-terminal myristoylation to increase the membrane affinity of the sensors

Next, we hypothesized that the likely transient and weak interactions between the candidate peptides and the GαGTPs generated at the plasma membrane are the underlying cause of the above defects. Although lipid modifications like myristoylation or farnesylation can target proteins to membranes, a lipid modification alone can only provide limited membrane affinity enhancement, which is usually traded off for the protein’s solubility and the membrane’s composition.^53,54^ When standalone plasma membrane targeting is desired, as seen in prenylated carboxy termini of small G proteins, positively charged residues are usually required for their plasma membrane targeting, as those residues form interactions with the negatively charged lipid headgroups of membrane phospholipids.^43^ Therefore, it is generally accepted that myristoylated or prenylated proteins require more than one membrane-targeting motif.^53-55^ For instance, Gαi is doubly lipidated and stays plasma membrane-bound even after dissociation from the heterotrimer, while the singly lipidated Gαs is mobilized to the cell interior upon liberation from Gβγ.^56-58^ To examine the feasibility of using lipid motifs to improve the GαGTP sensitivities of these candidate peptides, we tethered an N-terminal myristoylation sequence (MAS) to both the peptides and generated MAS-KB1753-Venus and MAS-GRK2-RH-Venus fusion constructs. To assess their functionality, we first expressed MAS-KB1753-Venus with α2AR-CFP in HeLa cells. Here, even before α2AR activation, MAS-KB1753-Venus was predominantly plasma membrane localized (Fig. 1g), making MAS-KB1753-Venus ineffective as a translocation-based GαiGTP sensor. The plasma membrane localization of MAS-KB1753-Venus was confirmed using its co-localization with G protein γ3 tagged with mCherry (Fig. 1g). Here, the line profile plot clearly shows that Venus and mCherry fluorescence peaks overlap, indicating their localization on the plasma membrane (Fig. 1g; the grey area indicates the plasma membrane region). Similarly, we examined MAS-GRK2-RH-Venus localization in HeLa cells also expressing GRPR and Gαq-CFP. Interestingly, unlike myristoylated KB1753-Venus, MAS-GRK2-RH-Venus was primarily cytosolic (Fig. 1h, 0 s). However, GRPR activation only induced a minor recruitment of MAS-GRK2-RH-Venus to the plasma membrane despite the exogenous Gαq expression (Fig. 1h, images of 90 s, 240 s, white arrows, and plot). In the absence of Gαq expression, even a minor recruitment of MAS-GRK2-RH-Venus was not observed (data not shown).

### Screening chaperones for myristoyl group to modulate MAS-KB1753-Venus localization

To reduce plasma membrane affinity, while increasing the solubility of the myristoylated sensor, we examined the feasibility of using a tethered chaperone. Here, we explored three potential lipid-interacting chaperone-like molecules: UNC119a, UNC119b,^59,60^ and PDEδ6^61^. To test them, we created MAS-KB1753-(chaperone)-Venus constructs. We expressed each construct alongside α2AR-CFP in HeLa cells. The KB1753 with UNC119a and UNC119b exhibited cytosolic distribution before receptor activation (Fig. 2a, cell images of 0 s). However, the PDEδ6 version remained predominantly localized to the plasma membrane, ruling out PDEδ6 as a myristoyl group chaperone (Fig. S2). Upon α2AR activation, we observed minor, but detectable plasma membrane recruitment of sensors with the added chaperones, UNC119a, and UNC119b, (Fig. 2a, cell images at 150 s, 300 s, and white arrows). To better understand the poor sensor performances observed here, we computationally analyzed the AlphaFold2 structures of these sensors.^62,63^ We first examined the conservation of each domain in these modeled modular AlphaFold2 structures with the available PDB structures. Moving forward, we selected modeled structures for subsequent in silico analyses that showed accurately predicted domains. These selected structures also exhibited the highest coverage scores (Supplementary Tables 1, 3, and 5).

**Figure 2.**
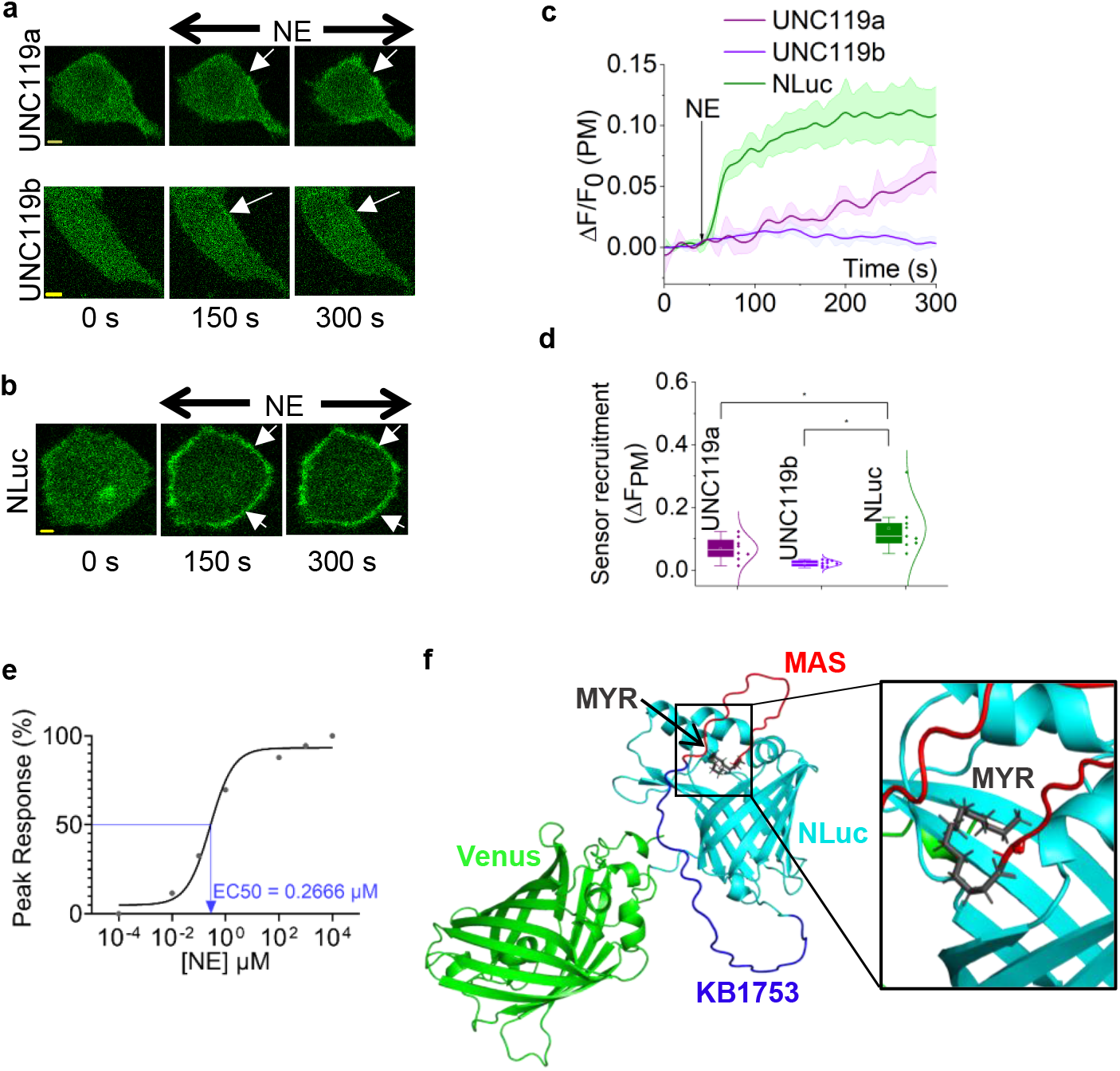
The EndoGi Tracker requires a chaperone for the myristoyl group for efficient sensor function. (a) (b) Hela cells expressing α2AR-CFP and MAS-KB1753-chaperone-Venus (chaperone: UNC119a, UNC119b, or NLuc) exhibit distinct plasma membrane recruitment patterns upon α2AR activation with 100 μM norepinephrine (white arrows). (c) The corresponding line plots show Venus fluorescence dynamics on the plasma membrane with UNC119a (violet plot, n = 9), UNC119b (purple, n = 11), or NLuc (green, n = 10). (d) The whisker box plot shows the EndoGi Tracker recruitment extents with UNC119a, UNC119b, or NLuc. MAS-KB1753-NLuc-Venus exhibits significantly higher plasma membrane recruitment extent compared to UNC119a and UNC119b. (e) Dose response curve of EndoGi Tracker recruitment to α2AR in response to a concentration gradient of norepinephrine (0.0001-10000 μM). EC50=0.2666 uM. (f) The Homology model of EndoGi Tracker (MAS-KB1753-NLuc-Venus) was generated using AlphaFold2 with the highest coverage score.The Schrödinger software was used to dock the myristoyl group to chaperone, Nluc. Docking hit with the best pose and highest docking score is shown here. The myristoyl group fits into the hydrophobic pocket of Nluc. Different regions of the protein are depicted in separate colors: MAS: red; KB1753: dark blue; Venus: green; myristoyl group: black; NLuc: cyan. Average curves were plotted using ‘n’ cells, n = number of cells. Error bars represent SEM (standard error of mean). The scale bar = 5 µm. Statistical comparisons were performed using One-way-ANOVA; p < 0.05, (*: population means are significantly different at 95% confidence level; ns: population means are not significantly different at 95% confidence level).

Next, we docked the myristoyl group to the protein models using GLIDE (Grid-based Ligand Docking with Energetic) tool in Schrödinger Maestro, and compared the docking scores and the Emodel. Docking score represents protein-ligand binding affinity, while Emodel scores provide a measure of combined protein-ligand interaction energy, docking score contribution, and ligand internal strain energy.^64,65^ The docking scores and Emodel values of all hits for PDEδ6, UNC119a, and UNC119b constructs are summarized in Supplementary Tables 2, 4, and 6. The data revealed distinct energy profiles for the myristoyl group binding. Interestingly, a few hits were observed for myristoyl-Venus interactions as well. However, as described above, since MAS-KB1753-Venus was plasma membrane-localized even before receptor activation, despite the presence of Venus, we discarded any hits exhibiting myristoyl-Venus interactions. Docking and Emodel scores of the most favorable myristoyl group binding site for all three chaperones, PDEδ6, UNC119a, and UNC119b were negative, indicating the tight binding of myristoyl anchor to the respective chaperone (Fig. S3, Supplementary Tables 2, 4, and 6). This explains the cytosolic localization and the lack of plasma membrane recruitment of the sensors either with UNC119a or UNC119b, upon GαiGTP generation. Likely, the tethered myristoyl group may not have access to the deeper binding pocket observed in PDEδ6, and may have prevented the cytosolic localization of this sensor candidate (Fig. S3). This is consistent with the ability of PDEδ6 to act as a prenyl group chaperone in KRAS, in which the lipid is tethered to the flexible C-terminus of the protein.^66^

The engineered luciferase, Nluc, shows that its core contains a binding site for the lipophilic luciferin substrate, coelentrazine, suggesting its potential as a lipid chaperone.^67^ We therefore generated MAS-KB1753-NLuc-Venus, as well, which exhibited a cytosolic distribution (Fig. 2b, s). Upon α2AR activation in HeLa cells, this sensor exhibited robust plasma membrane recruitment (Fig. 2b, 150 and 300 s, and Supplementary Movie 1), suggesting that Nluc provided sufficient chaperone activity, thereby allowing plasma membrane recruitment of the sensor using the additional affinity provided by GαiGTP. This observation also helped us to propose that the MAS-KB1753-Nluc used in the BRET-based detection of GDP-to-GTP exchange on the exogenous Gαi (in the ONE-GO biosensor)^27^ was feasible because Nluc, in this context, too, acts as a chaperone for the myristoyl group. Image analysis clearly showed that Nluc outperformed the other chaperones examined (Fig. 2c and 2d, plots, and Supplementary Table 11). Therefore, MAS-KB1753-NLuc-Venus was named the EndoGi Tracker for subsequent experiments. We next examined α2AR-norepinephrine dose response in HeLa cells using our novel EndoGi tracker. The dose-response curve showed an EC50 value of 0.27 µM norepinephrine, and the peak activity was reached at ∼100 µM (Fig. 2e and S5a: cell images). Next, we generated AlphaFold structures for the EndoGi Tracker using the same procedure mentioned above (Supplementary Table 7). The docking score of the myristoyl group with the AlphaFold structure of the EndoGi Tracker (Fig. 2f) was positive compared to the previously tested chaperones, indicating a more transient interaction with Nluc (Supplementary Table 8). We therefore propose that these transient myristoyl-Nluc interactions underlie the robust EndoGi tracker plasma membrane recruitment upon the additional affinity conferred by GαiGTP generation.

### The EndoGi Tracker is a powerful tool for detecting subcellular opioid receptor activity

The detection of dynamic signaling from Gi-coupled GPCRs has been a significant limitation in pharmacology and GPCR deorphanization research. Pain perception is one of the major physiological processes regulated by GPCRs, including opioid, cannabinoid, serotonergic, and glutamate receptors, and the majority of these nociceptors are Gi/o-coupled.^68-73^ Distinct localizations of these receptors in different cell types can determine the functional outcomes of their agonists.^72,74^ For instance, receptors, including kappa opioid and cannabinoid receptors, elicit distinct signaling outcomes depending on their subcellular localization.^75,76^ However, to our knowledge, how the nociceptors activate endogenous G protein heterotrimers in single cells has not been reported, likely due to the unavailability of capable probes. Therefore, we next employed EndoGi Tracker to detect opioid receptor signaling in real time in live cells. First, we expressed µ-opioid Receptor (MOR) and EndoGi Tracker in HeLa cells. Before receptor activation, the EndoGi Tracker was cytosolic (Fig. 3a, 0 s), and upon receptor activation with 1 nM Fentanyl, the sensor translocated to the plasma membrane, indicating robust GαiGTP generation with a half-time (t_1/2_) of 12.43 ± 3.22 s (Fig. 3a, 100 and 300 s, white arrows, and plot). Next, the Receptor was inhibited by adding 2 nM naloxone (MOR competitive antagonist), which showed a near-complete recovery of the EndoGi Tracker back to the cytosol with a t_1/2_ of 46.34 ± 12.37 s (Fig. 3a, 600 s, yellow arrow, and plot). Half-time distributions for activation and inactivation of MOR are shown in Fig. 3b. This ∼4-fold slower reduction of Gαi-GTP (Fig. 3b and Supplementary Table 12) is consistent with the involvement of GTPase activity in the signaling cessation of heterotrimeric G proteins and indicates that EndoGi Tracker is an effective sensor for detecting on-off signaling kinetics downstream of GPCRs. To ensure that EndoGi Tracker is compatible with other nociceptors, we examined GαiGTP dynamics in HeLa cells upon Cannabinoid 1 receptor (CB1R) activation. Here, we expressed CB1R and EndoGi Tracker in HeLa cells. Upon CB1R activation with the synthetic ligand, WIN-55,212-2 (50 nM), EndoGi Tracker robustly recruited to the plasma membrane with a t_1/2_ = 16.27 ± 2.58 s (Fig. 3c, 240 and 480 s, white arrows, and plot). With 50 nM CB1R antagonist, AM251, we observed a full recovery of the EndoGi Tracker back to the cytosol with a t_1/2=_144.79 ± 36.49 s (Fig. 3c, 720 and 960 s, yellow arrows, and plot). The half-time distributions for activation and inactivation of CB1R are shown in Fig. 3d. Again, we observed significantly slower return (∼9-fold) of the EndoGi Tracker upon CB1R inactivation (Fig. 3d and Supplementary Table 13) This data collectively indicate that our EndoGi Tracker can be used as a universal sensor that detects GαiGTP generation in real-time, enabling the explicit probing of distinct nociceptor pathways to obtain dynamic molecular pictures unique to ligands and receptors, and decode distinct pharmacological properties likely to be crucial in developing novel therapeutic strategies for pain management with minimal deleterious effects.

**Figure 3.**
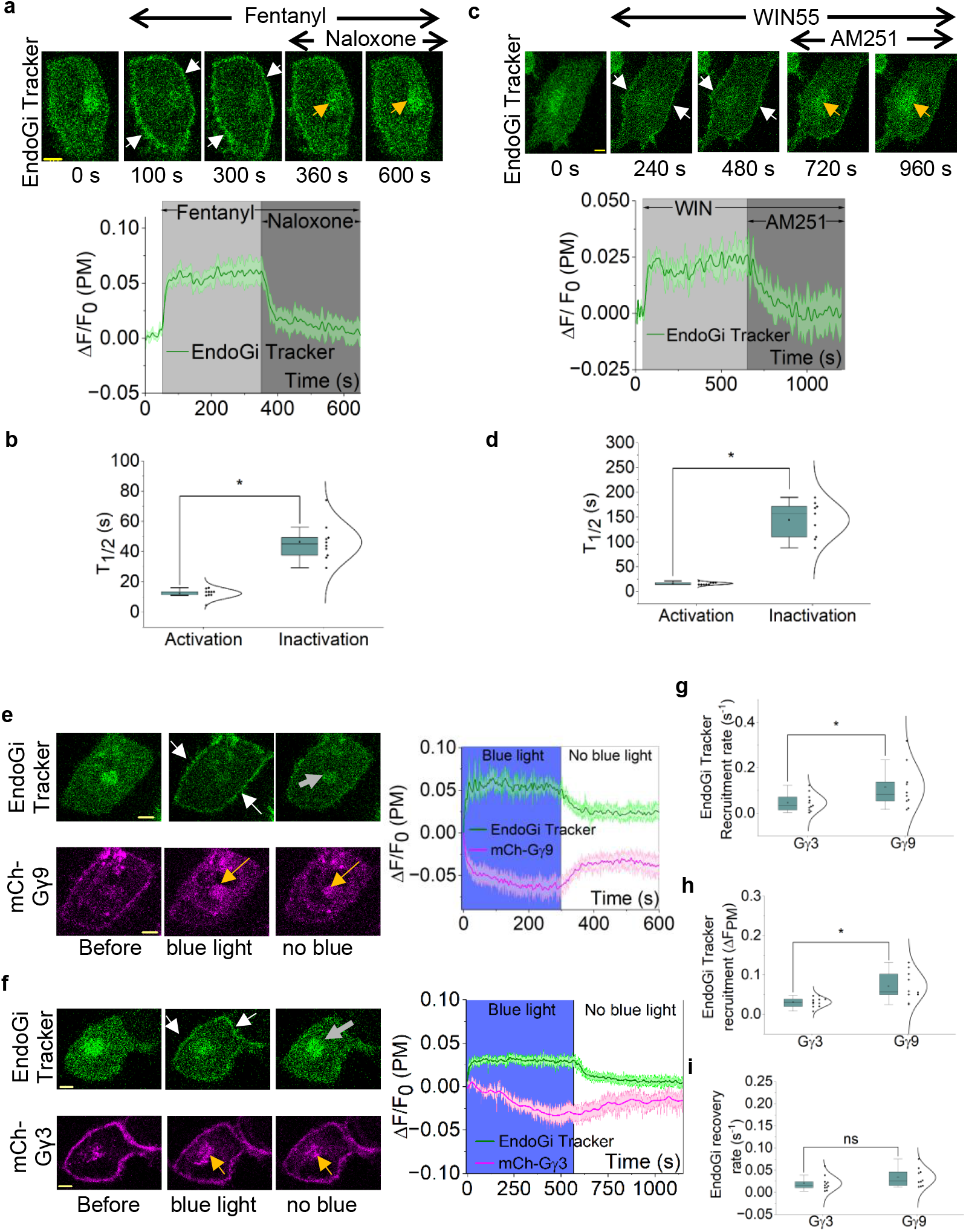
EndoGi Tracker is a powerful tool for detecting subcellular GPCR-G protein on-off kinetics. (a) Fentanyl induced robust EndoGi Tracker recruitment to the plasma membrane in HeLa cells expressing µ-opioid Receptor (MOR)-mCh with EndoGi Tracker (white arrows). Plasma membrane-bound EndoGi Tracker showed near-complete recovery upon receptor inactivation with naloxone (yellow arrows). The corresponding plot depicts intracellular movement of the sensor upon MOR-activation-deactivation signaling (n = 10). (b) The box plot shows the EndoGi Tracker half-times (T_1/2_) for recruitment and recovery upon MOR activation and inactivation, respectively. (c) HeLa cells expressing cannabinoid receptor 1 (CB1R) with EndoGi Tracker showed robust sensor recruitment to the plasma membrane (white arrows) upon CB1R activation with WIN-55,212-2. Addition of the inhibitor AM251 resulted in near-complete recovery of the EndoGi Tracker to the cytosol (yellow arrows). The plot shows EndoGi Tracker dynamics on the plasma membrane (n = 9). (d) The box plot shows the EndoGi Tracker recruitment and recovery half times (T_1/2_) upon CB1R activation and inactivation, respectively. (e) (f) HeLa cells expressing blue opsin-mTq, EndoGi Tracker with either mCh-Gγ9 or mCh-Gγ3 showed robust sensor recruitment with blue light activation (white arrows), and the sensor recovered back to the cytosol upon removal of blue light (grey arrows). Both mCh-Gγ9 and mCh-Gγ3 exhibited translocation from the plasma membrane to internal membranes upon opsin activation with blue light and recovery back to the plasma membrane upon blue light removal (yellow arrows). (g) (h) EndoGi Tracker recruitment extent and rate were significantly higher in Gγ9-expressing cells over Gγ3-expressing cells. (i) Interestingly, EndoGi Tracker recovery rates (upon receptor inactivation) did not show a significant difference between the two conditions. Average curves were plotted using ‘n’ cells, n = number of cells. Error bars represent SEM (standard error of mean). The scale bar = 5 µm. Statistical comparisons were performed using One-way-ANOVA; p < 0.05, (*: population means are significantly different at 95% confidence level; ns: population means are not significantly different at 95% confidence level).

### EndoGi Tracker shows that Gi-pathway-mediated heterotrimer signaling is Gγ-subtype-dependent

To our knowledge, there are no sensors available to detect endogenous G protein heterotrimer activation (Gα and βγ dissociation) and inactivation (heterotrimer re-assembly) in single cells. Therefore, we employed the EndoGi tracker to examine the regulation of endogenous Gαi heterotrimer activation. We have previously shown that the activation efficacy of Gβγ-mediated downstream effectors, such as PI3Kγ and PLCβ, is dependent on the Gγ subtype in the heterodimer.^31,44^ Based on that, we also wanted to test the hypothesis that, since Gγ controls the membrane affinity and lateral diffusion of Gβγ, it also controls GPCR-heterotrimer interactions in a Gγ-type-dependent manner. To test this, we expressed blue opsin, a light-activatable Gi-coupled GPCR, tagged with mTurquoise and EndoGi tracker in HeLa cells also expressing either mCh-Gγ3 or mCh-Gγ9. We selected Gγ3 and Gγ9 because they exhibit the highest and the lowest membrane affinities, respectively.^29,43^ We used blue opsin as the Gi-coupled GPCR to precisely turn the opsin on and off at user-defined time points.^77,78^ After the addition of 10 µM 11-cis-retinal (to generate optically activatable blue opsin), we exposed cells to blue light to activate the receptor while imaging Venus and mCherry to examine the EndoGi Tracker and Gγ, and then terminated the blue light to deactivate the receptor. Here, upon blue light, we observed robust plasma membrane recruitment of the EndoGi Tracker. Although we observed a robust mCh-Gγ9 translocation from the plasma membrane to internal membranes, Gγ3 exhibited a slower response with a lower extent (Fig. 3e and f, blue light, white and yellow arrows, and plots). Blue light termination (cessation of mTurquoise imaging) resulted in the recovery of the EndoGi Tracker and the reversal of Gγ back to the plasma membrane (Fig. 3e and f; no blue light, grey and yellow arrows, respectively, and plots). Next, we calculated the dynamics of sensor recruitment and recovery over time. The EndoGi Tracker recruitment rate and extent were significantly higher (∼2.5-fold) in Gγ9-expressing cells (Fig. 3g, h, and Supplementary Tables 14 and 15), while Gγ subtype did not influence the EndoGi Tracker reverse upon blue light termination (Fig. 3i and Supplementary Table 16). These data indicate that Gγ subtype influences G protein activation and heterotrimer dissociation upon GPCR activation. Since Gγ9 has the lowest membrane affinity, it may allow Gγ9 heterotrimers to laterally diffuse faster along the plasma membrane, while enabling better shuttling between membranes than those of Gγ3 heterotrimers. This likely helps Gγ9 heterotrimers readily dissociate and efficiently generate GαGTP. However, as expected, the GαGTP hydrolysis upon signaling termination was not Gγ-type dependent, as this process is Gβγ independent. This observation is significant because distinct cell and tissue types exhibit different predominant, 2-3 Gγ subtypes in a given cell type, and therefore, for the same ligand and receptor, the onset of heterotrimer activation kinetics and G protein-mediated downstream signaling can vary depending on the Gγ expression profile.^31,43^

### Engineering of EndoGq Tracker and examination of endogenous Gαq-GTP dynamics

Although the GαqGTP sensor candidate, MAS-GRK2-RH-Venus exhibited proper cytosolic localization (Fig. 1h, 0 s), since it showed negligible GαqGTP sensitivity, we first computationally examined possible reasons for this behavior. Here, we employed an approach similar to the above (Fig. 2f and S3) and generated an AlphaFold2 model of the modular protein and examined the structural conservation of each module relative to its corresponding PDB structure (2BCJ). None of the obtained hits maintained the structural integrity of the GRK2-RH domain (Fig. S4a). We therefore hypothesized that the GRK2-RH orientation in MAS-GRK2-RH-Venus reduces its GαqGTP binding ability. Thus, we revised the modular architecture of the sensor to create MAS-Venus-GRK2-RH, to improve the spatial flexibility required for structural integrity, which may also be necessary for GRK2-RH—GαqGTP interactions to occur. Interestingly, in all AlphaFold2 MAS-Venus-GRK2-RH structure hits, the GRK2-RH structure was preserved (Fig. S4b).

The revised sensor, MAS-Venus-GRK2-RH, not only exhibited cytosolic localization in HeLa cells (Fig. 4a, 0s), but it also exhibited robust translocation to the plasma membrane upon GRPR activation with 1 µM bombesin (Fig. 4a, 90 and 240 s, white arrows, and plot). Here, we exogenously expressed Gαq since HeLa cells endogenously express only a limited Gαq.^30^ HeLa cells in a similar experiment without Gαq expression also exhibited a lower, however, clear sensor recruitment to the plasma membrane (Fig. 4b, 180 s, 300 s, and white arrows). This data clearly indicated that the sensor is able to detect endogenous GαqGTP in cells, even with low abundance of endogenous Gαq. Therefore, hereafter, the MAS-Venus-GRK2-RH sensor is referred to as the EndoGq Tracker. When comparing the EndoGq Tracker recruitment kinetics, Gq overexpressed (+Gαq) cells showed a significantly greater extent (∼4-fold) and faster rate (∼9-fold) (Fig. 4c and 4d, Supplementary Tables 17 and 18) compared to that of the native (-Gαq) condition. However, with endogenous Gαq, the EndoGq Tracker exhibited significant recovery (∽71.85%) over time (Fig. 4b, 600 s, yellow arrow, 4e, and Supplementary Table 19).

**Figure 4.**
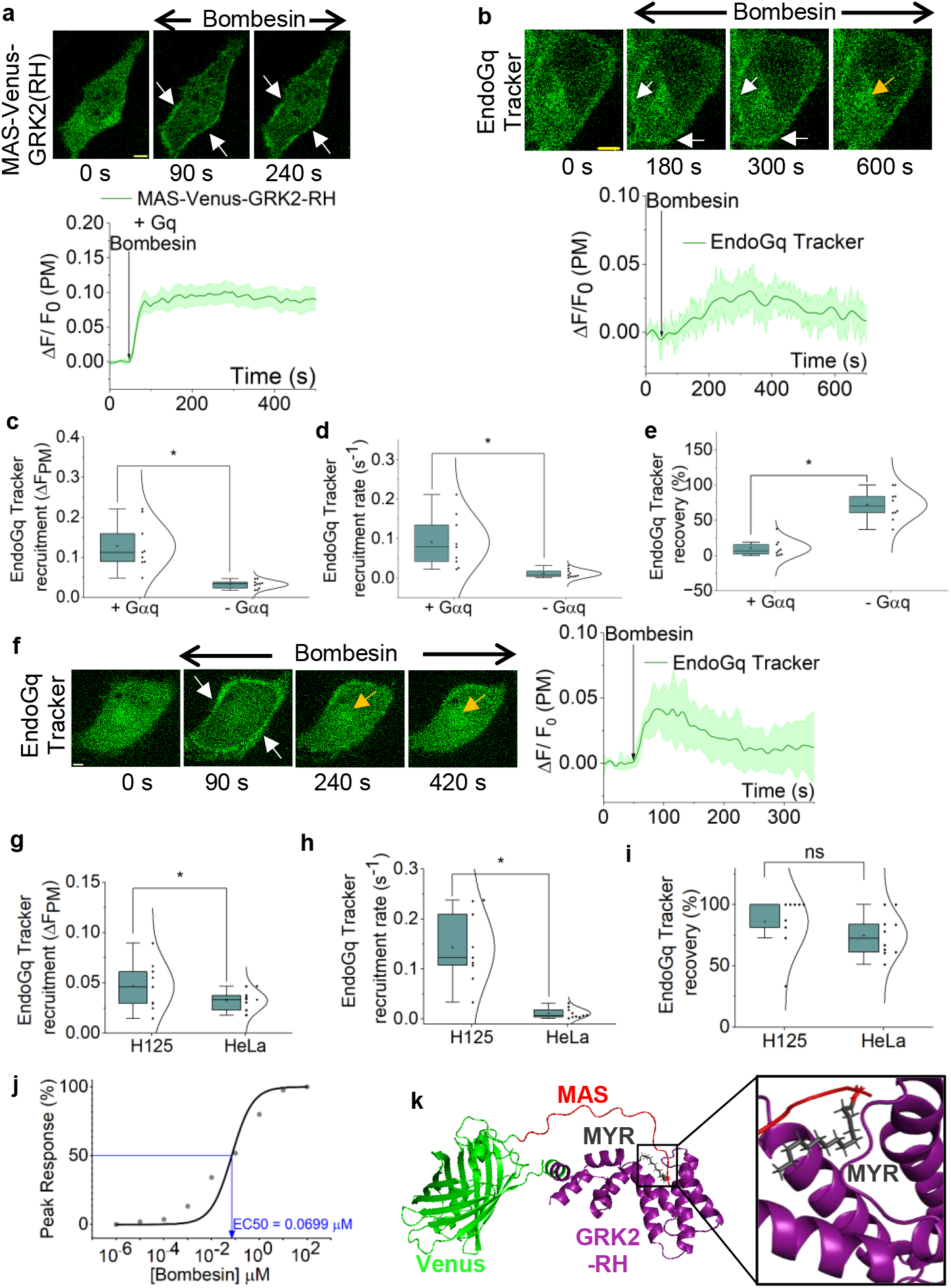
Cytosolic MAS-Venus-GRK2-RH efficiently detects GαqGTP generation and translocates to the plasma membrane. (a) Hela cells expressing GRPR, MAS-Venus-GRK2-RH, and Gαq-CFP exhibit robust plasma membrane recruitment upon GRPR activation with 1 μM bombesin (white arrows). The corresponding plot shows MAS-Venus-GRK2-RH dynamics on the plasma membrane (n = 9). (b) A minor but detectable recruitment of MAS-Venus-GRK2-RH was observed upon GRPR activation in HeLa cells lacking exogenous Gαq expression (white arrows). Interestingly, MAS-Venus-GRK2-RH recovered to the cytosol (600 s: yellow arrow). The corresponding plot shows the MAS-Venus-GRK2-RH profile on the plasma membrane (n = 11). (c) (d) The EndoGq Tracker (MAS-Venus-GRK2-RH) recruitment extent and rate were significantly higher in Gαq-expressing HeLa cells compared to cells without exogenous Gαq expression. (e) The subsequent EndoGq Tracker recovery was significantly higher in cells with endogenous Gαq. (f) The EndoGq Tracker exhibited robust plasma membrane recruitment upon GRPR activation in NCI-H125 cells (white arrows) expressing GRPR alongside the sensor and subsequently recovered to the cytosol (yellow arrows). The corresponding plot shows the EndoGq Tracker dynamics on the plasma membrane (n = 9). (g) (h) The EndoGq Tracker recruitment extent and rate were significantly higher in NCI-H125 cells compared to HeLa cells. (i) However, the EndoGq Tracker recovery extent did not show any significant difference between NCI-H125 cells and HeLa cells. (j) Dose response curve of EndoGq Tracker recruitment to GRPR in response to a concentration gradient of bomesin (0.001 nM – 100 μM) in NCI-H125 cells (EC50=0.0699 nM). (k) The structure of EndoGq Tracker (MAS-Venus-GRK2-RH) modeled from AlphaFold2 modeling software with the highest coverage score. The Schrödinger software was used to dock the myristoyl group to the modeled protein. Docking hit with the best pose and highest docking score is shown here. The myristoyl group fits into a hydrophobic pocket in GRK2-RH. Different regions of the protein are depicted in separate colors: MAS: red; Venus: green; myristoyl group: black; GRK2-RH: purple. Average curves were plotted using ‘n’ cells, n = number of cells. Error bars represent SEM (standard error of mean). The scale bar = 5 µm. Statistical comparisons were performed using One-way-ANOVA; p < 0.05, (*: population means are significantly different at 95% confidence level; ns: population means are not significantly different at 95% confidence level).

To examine whether the observed lower (Fig. 4c), as well as slower (Fig. 4d) sensor recruitment in HeLa cells with endogenous Gαq (Fig. 4b) is indicative of Gq expression, we also probed EndoGq Tracker behavior in NCI-H125 cells, since this cell line has been reported to have an upregulated endogenous Gαq expression.^30^ In NCI-H125 cells expressing GRPR alongside EndoGq Tracker, upon activation of GRPR with 1 µM bombesin, we observed robust recruitment of the EndoGq Tracker to the plasma membrane (Fig. 4f, 90 s, white arrows, and plot). However, the sensor exhibited a near-complete recovery back to the cytosol (Fig. 4f, 240 and 420 s, yellow arrows, and plot). Supplementary Movie 2 exhibits real-time EndoGq recruitment and subsequent recovery in a NCI-H125 cell. Next, we calculated the extent of EndoGq Tracker recruitment extent, the recruitment rate, and the subsequent recovery extent in H125 and HeLa cells lacking Gαq expression (-Gαq condition). Here, we observed ∽1.5-fold higher sensor recruitment (Fig. 4g and Supplementary Table 20) and ∽12.5-fold faster sensor recruitment (Fig. 3h and Supplementary Table 21) in H125 cells than in HeLa cells. This observation also underscores the utility of this sensor for estimating relative endogenous Gαq expression levels. However, the sensor recovery did not show any significant difference between NCI-H125 and HeLa cells with endogenous Gαq (Fig. 4i and Supplementary Table 22). This observed sensor recovery is not only consistent with our previously reported PIP2 hydrolysis adaptation in HeLa cells with endogenous Gαq,^44^ but also appears to be conserved when Gαq is not exogenously introduced into NCI-H125 cells. We next utilized our newly engineered EndoGq Tracker to investigate the dose-response relationship of GRPR-mediated GαGTP generation in NCI-H125 cells. Here, we observed an EC50 of 0.07 µM and peak activity at concentrations above 1 µM of bombesin (Fig. 4j and S5b: cell images). Next, we docked the myristoyl group to the AlphaFold structure of EndoGq tracker with the highest coverage score (Supplementary Table 9) in a manner similar to that previously described. Interestingly, in this global docking, after excluding hits involving Venus, we identified one hit showing myristoyl group interaction with GRK2-RH (Fig. 4k and Supplementary Table 10), indicating the GRK2-RH domain itself likely acts as a chaperone for the myristoyl group.

We also examined whether the EndoGq tracker’s response is universal across all Gq-coupled GPCRs. We expressed muscarinic 3 (M3) receptor with the EndoGq tracker in NCI-H125 cells. Upon activation of M3 receptor with 10 µM Carbachol, we observed robust EndoGq tracker recruitment to the plasma membrane (Fig. 5a, 110 s, white arrows, and plot) and subsequent recovery (Fig. 5a, 240 and 420 s, yellow arrows, and plot), similar to GRPR, suggesting that sensor recovery is common to other Gq receptors as well.

**Figure 5.**
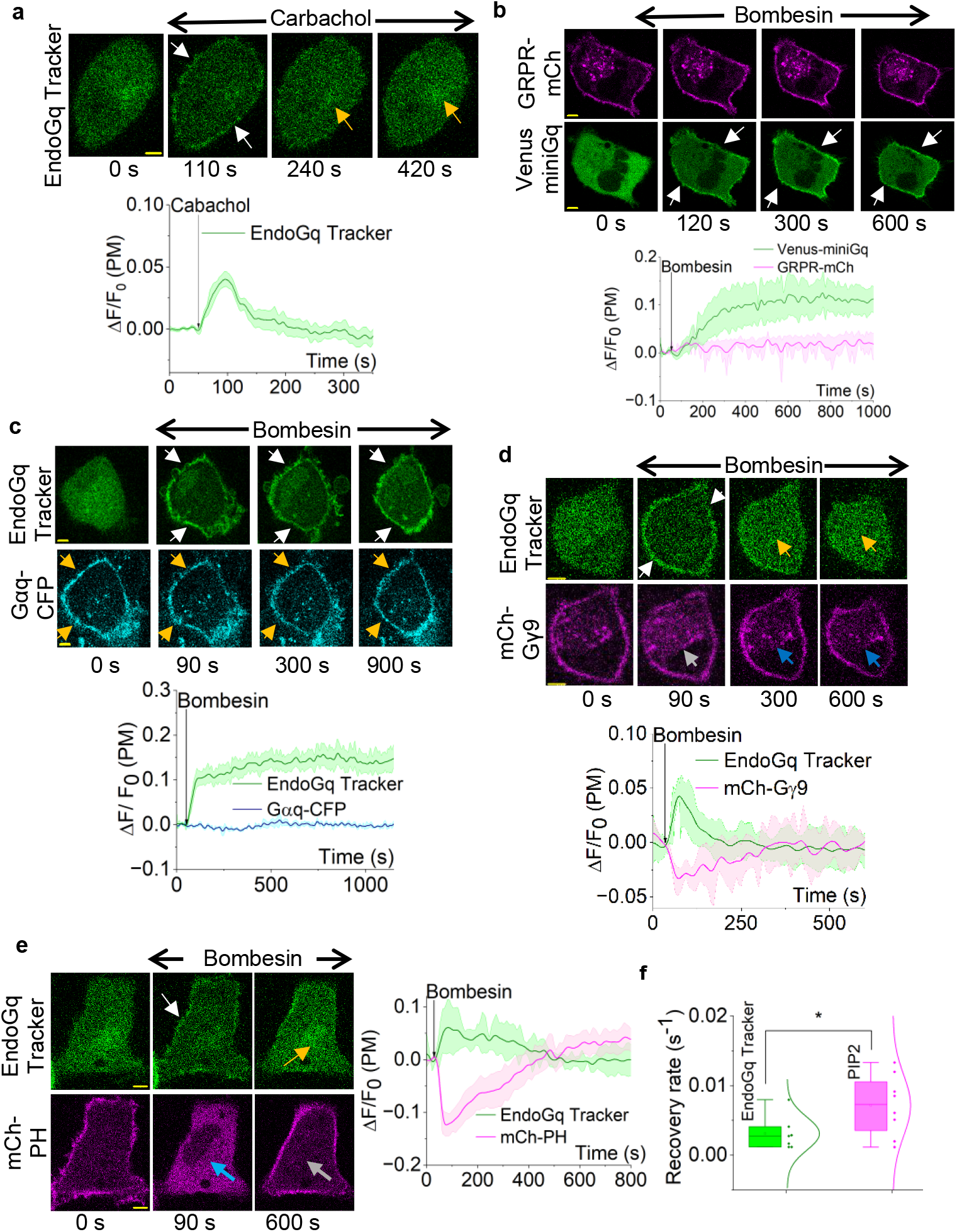
EndoGq Tracker reveals non-canonical cellular mechanisms to overcome Gq pathway overstimulation. (a) Upon muscarinic 3 receptor (M3R) activation with 10 µM Carbachol, the EndoGq Tracker showed plasma membrane recruitment in NCI-H125 cells (white arrows). Plasma membrane-bound sensor eventually returned to the cytosol despite the continuous presence of stimulus (yellow arrows). The plot shows EndoGq Tracker dynamics on the plasma membrane (n = 10) (b) NCI-H125 cells expressing GRPR-mCh and Venus-miniGq exhibited robust miniGq recruitment to the plasma membrane upon GRPR activation, while GRPR stayed on the plasma membrane throughout the time course of the experiment. The plot shows the mean fluorescence intensity profiles of Venus-miniGq (green) and GRPR-mCh (magenta) (n = 9). (c) NCI-H125 cells expressing GRPR, Gαq-CFP, and EndoGq Tracker exhibit robust sensor recruitment upon GRPR activation. Interestingly, exogenous Gαq expression resulted in little to no recovery of the EndoGq Tracker to the cytosol and continued to stay on the plasma membrane (white arrows). Gαq-CFP stayed on the plasma membrane throughout the experiment (yellow arrows). The plot shows the fluorescence intensity profiles of EndoGq Tracker (green) and Gαq-CFP (blue) (n = 9). (d) NCI-H125 cells expressing GRPR, EndoGq Tracker, and mCh-Gγ9 exhibit simultaneous robust EndoGq Tracker recruitment (white arrows) and Gγ9 translocation (grey arrows), upon GRPR activation. Interestingly, both the EndoGq Tracker (yellow arrows) and Gγ9 (blue arrows) recovered, indicating heterotrimer reformation. The corresponding plot shows the EndoGq Tracker (green) and mCh-Gγ9 (magenta) dynamics on the plasma membrane (n = 9). (e) NCI-H125 cells expressing GRPR, EndoGq Tracker, and PIP2 sensor, mCh-PH exhibit robust EndoGq Tracker recruitment to the plasma membrane (white arrows) and PIP2 hydrolysis (indicated by the PIP2 sensor moving to the cytosol from the plasma membrane, blue arrow), simultaneously upon GRPR activation. Both the EndoGq Tracker (yellow arrows) and mCh-PH (grey arrows) recovered back to the cytosol and the plasma membrane, respectively. The corresponding plot shows the dynamics of the EndoGq Tracker (green) and mCh-PH (magenta) (n = 9). (f) The rate of EndoGq Tracker recovery to the cytosol was significantly slower than the PIP2 sensor recovery to the plasma membrane. Average curves were plotted using ‘n’ cells, n = number of cells. Error bars represent SEM (standard error of mean). Statistical comparisons were performed using one-way ANOVA; p < 0.05, (*: population means are significantly different at 95% confidence level). The scale bar = 5 µm.

Since the EndoGq tracker detects GαqGTP, the sensor recovery observed could stem from several factors, including GPCR desensitization, translocation of generated GαqGTP to the cytosol over time, rapid GTP hydrolysis on Gαq, reduced heterotrimer activation by the GPCR, or the sensor competing with GαqGTP effectors. We therefore explored the molecular details underlying this phenomenon.

### Real-time analysis of Gαq heterotrimer activation and downstream signaling dynamics

Initially, we hypothesized that Gαq heterotrimer activation diminishes over time due to GPCR desensitization and internalization. To assess this, we expressed GRPR-mCh and Venus-miniGq in HeLa cells, and upon activation of GRPR with 1 µM bombesin, cytosolic miniGq translocated to the plasma membrane, indicating robust GPCR activity (Fig. 5b, white arrows, and plot). Interestingly, miniGq remained translocated to the plasma membrane for the duration of the experiment (∽20 min), indicating that GRPR remained active. Also, GRPR did not show any detectable internalization (Fig. 5b, cell images: GRPR-mCh, and magenta plot). This suggested that the EndoGq recovery is not likely due to GPCR desensitization.

Next, we tested whether GαqGTP translocates to the cytosol or if the interaction between GαqGTP and the EndoGq Tracker is transient and superseded by additional mechanisms, such as an effector interaction with GαqGTP. Therefore, we first examined whether the observed recovery is due to GαqGTP translocation to the cytosol, similar to the reported Gαs translocation.^58,79^ We expressed GRPR and EndoGq tracker with Gαq-CFP in NCI-H125 cells. As expected, the EndoGq tracker was localized to the cytosol (Fig. 5c, EndoGq tracker: 0 s), while Gαq-CFP exhibited a plasma membrane localization (Fig. 5c, Gαq-CFP: 0 s, yellow arrows). Upon receptor activation with 1 µM bombesin, we observed rapid EndoGq tracker recruitment to the plasma membrane (Fig. 5c, 90 s, 300 s, and 900 s, white arrows, and plot), while Gαq-CFP remained plasma membrane bound (Fig. 5c, 90, 300, and 900 s, yellow arrows, and blue plot). This suggested that EndoGq tracker recovery is not due to Gαq translocation away from the plasma membrane. Additionally, exogenous Gαq expression eliminated the sensor recovery observed with endogenous Gαq (Fig. 4f, and plot).

We also tested the hypothesis that the reduction of GαqGTP underscores the sensor recovery observed. Since Gβγ is liberated upon GαqGTP generation, we here examined whether sensor recovery is also associated with a reduction in free Gβγ. In NCI-H125 cells expressing GRPR, EndoGq tracker, and mCh-Gγ9, we performed the Gγ9 translocation assay to measure heterotrimer activation. We have shown that this assay can be used to measure Gq-GPCR activation in cells like NCI-H125 because of their relatively higher Gαq expression.^30^ Before GRPR activation, the EndoGq tracker was cytosolic, while mCh-Gγ9 was plasma membrane-bound (Fig. 5d, cell images of 0 min). Bombesin addition induced EndoGq tracker recruitment to the plasma membrane (Fig. 5d, 90 s, white arrows and plot) and Gγ9 translocation to the internal membranes (Fig. 5d, 90 s, grey arrow, and plot) simultaneously. Continuous imaging showed that the sensor recovered back to the cytosol, while Gγ9 returned to the plasma membrane (Fig. 5d, cell images of 300 s, 600s, and plot; yellow arrows: EndoGq tracker recovery; blue arrows: Gγ9 dynamics on the internal membranes). These observations can result from a transient increase in GαqGTP concentration at the plasma membrane upon Gq-GPCR activation (EndoGq tracker recruitment to the plasma membrane), together with a significant reduction in GαqGTP and an increase in GαqGDP. The observed Gγ9 recovery to the plasma membrane indicates heterotrimer reformation with GαqGDP. To ensure that EndoGq tracker expression does not interfere with or modulate Gβγ signaling, we examined GRPR-induced Gγ9 translocation with and without EndoGq Tracker expression in NCI-H125 cells. Here, we observed robust Gγ9 translocation and subsequent recovery independent of EndoGq tracker expression (Fig. S6a and S6b, and Supplementary Table 24), indicating that EndoGq tracker expression does not interfere with the Gαq pathway-induced Gγ9 translocation and recovery. These observations also suggest that the EndoGq Tracker-Gαq interactions do not significantly interfere with the Gq signaling adaptation and subsequent heterotrimer reformation.

In addition, we examined whether GαqGTP hydrolysis is reflected in the attenuation of its downstream signaling. GαqGTP is a potent activator of PLCβ, which hydrolyzes phosphatidylinositol 4,5-bisphosphate (PIP2) to diacylglycerol (DAG) and inositol triphosphate (IP3).^33^ Here, we expressed GRPR, EndoGq tracker, and mCh-PH (PIP2 sensor) in NCI-H125 cells. Before GRPR activation, mCh-PH was localized to the plasma membrane, as PIP2 is a plasma membrane-bound phospholipid, while the EndoGq tracker was cytosolic (Fig. 5e, 0 s). Upon GRPR activation with 1 µM bombesin, we observed robust PIP2 hydrolysis (Fig. 5e, 90 s, blue arrow, and plot) and EndoGq tracker recruitment to the plasma membrane (Fig. 5e, 90 s, white arrow, and plot). However, a significant, rapid PIP2 recovery was observed, as indicated by the reverse translocation of mCh-PH to the plasma membrane (Fig. 5e, 600 s, grey arrow) and by the sensor returning to the cytosol (Fig. 5e, 600 s, yellow arrow). However, a comparison of kinetics showed that the EndoGq tracker recovery is significantly slower (∼2-fold) than the PIP2 hydrolysis attenuation (Fig. 5f and Supplementary Table 23). This indicated that, in addition to the GαqGTP hydrolysis, the PIP2 recovery is likely governed by the molecular process we previously discovered, in which the translocation of generated Gβγ away from the plasma membrane underlies the attenuation of PIP2 hydrolysis.^44^

Since exogenous expression of Gαq-CFP inhibited EndoGq Tracker recovery (as in Fig. 5c), we examined whether this is reflected in downstream signaling as well. We explored Gγ9 translocation and PIP2 hydrolysis in Gαq-expressed NCI-H125 cells. Here, we observed that the attenuations of Gγ9 translocation and PIP2 hydrolysis (Fig. S7 and plot) were also abolished. Overall, this data shows that upon Gq pathway activation, GαqGTP undergoes a mechanism that results in signaling adaptation, regenerating the Gq heterotrimer. However, exogenous Gαq expression appeared to overcome this intrinsic adaptation mechanism, suggesting distinct signaling in cells with limited Gαq expression compared to those with elevated Gαq expression. One possibility is that, over time, efficacy heterotrimer activation by receptors is attenuated, while the higher heterotrimer density may still allow considerable GαGTP generation under these suboptimal conditions. Nevertheless, further investigation is required to determine the exact molecular details underlying the Gq signaling adaptation.

Moreover, to ensure that the results observed with our newly engineered sensors are not artifacts of photobleaching, receptor expression, or the cell types used, we performed negative control experiments in which both EndoGi Tracker and EndoGq Tracker, and corresponding GPCRs were expressed in HeLa and NCI-H125 cells, respectively, and only exposed to the vehicle solution of the respective ligand. No sensor recruitment was observed (Fig. S9a, S9b, and plots).

### EndoGαi and EndoGαqTrackers allow subcellular GαGTP detection

To examine the feasibility of using the new sensors to detect GαGTP generation upon localized GPCR activation in single cells, we expressed blue opsin with the mCh-tagged EndoGi Tracker in HeLa cells. While imaging cells for mCherry fluorescence to visualize the sensor, we exposed a confined region of the plasma membrane to 488 nm light (Fig. 6a, blue box). The localized blue opsin activation induced asymmetric GαiGTP generation on one side of the cell, as indicated by the sensor recruitment (Fig. 6a, 50 s, and white arrow, and line plot: Side A). Afterward, upon switching the 488 nm light stimulus to the opposite side, we were able to change the direction of GαiGTP generation and sensor recruitment (Fig. 6a, 165 s, grey arrow, and plot: Side B). Likewise, we were able to reversibly activate blue opsin on opposite sides by simply switching the 488 nm light stimulus (Fig. 6a, 290 s, and white arrow, and line plot: Side A). Next, we exposed the whole cell to blue light, which induced EndoGi Tracker recruitment to the plasma membrane globally (Fig. 6a, 375 s; yellow arrows and line plot). Real-time EndoGi Tracker dynamics with localized blue opsin activation is shown in Supplementary Movie 3. This data clearly indicates the utility of the EndoGi Tracker in detecting GαiGTP dynamics in subcellular regions. Bistable opsins are valuable optogenetic tools due to their potential in in vivo applications.^80,81^ Therefore, we examined EndoGi Tracker subcellular activity with the Gi-coupled bistable opsin, Lamprey parapinopsin.^81,82^ Similar to blue opsin, we observed robust, localized EndoGi Tracker recruitment to localized blue light (Fig. S8), indicating the potential utility of the sensor for acquiring subcellular signaling information.

**Figure 6.**
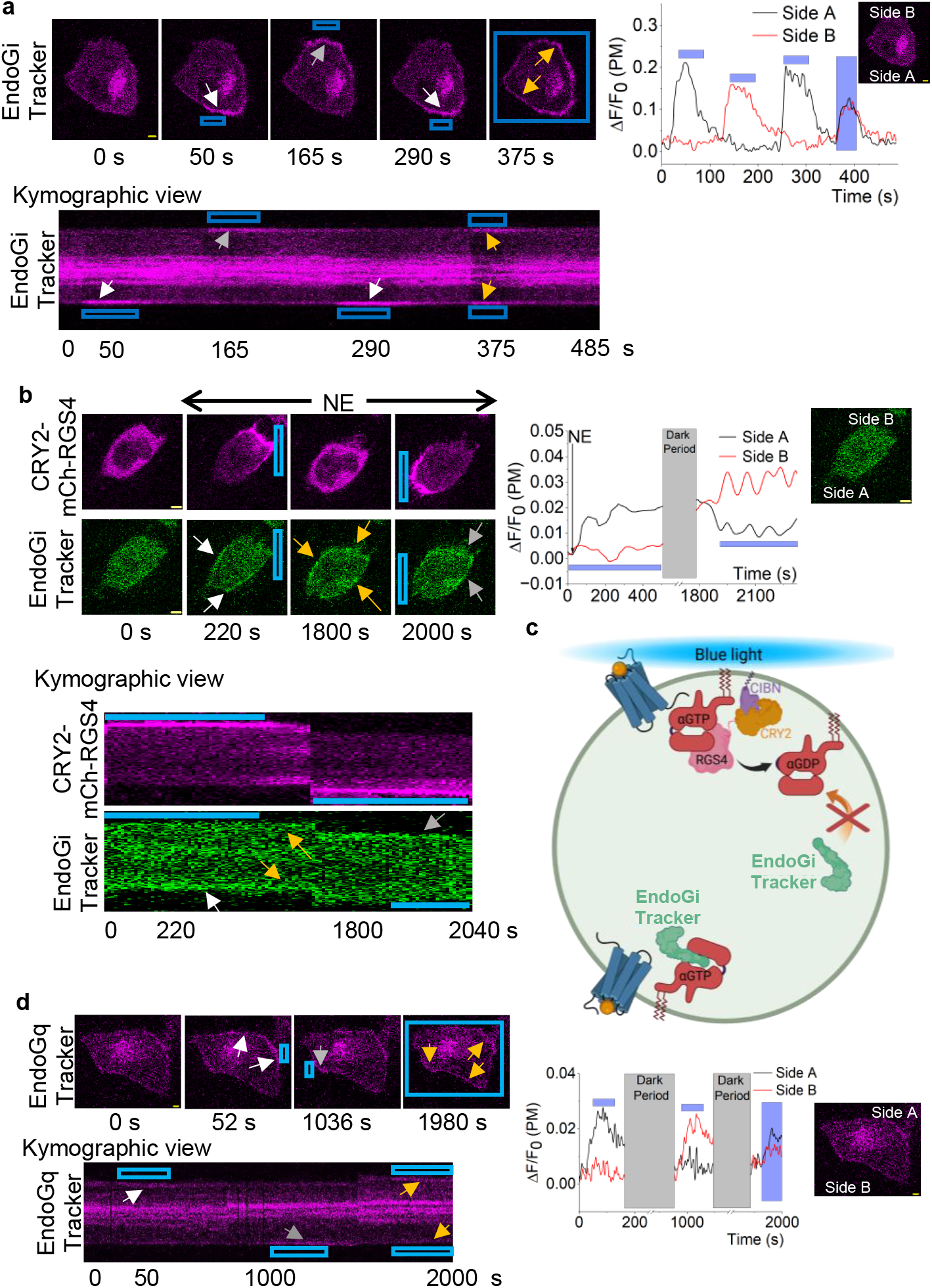
EndoGi Tracker and EndoGq Tracker detect GαGTP in subcellular regions. (a) HeLa cells expressing blue opsin and the EndoGi Tracker exhibit sensor recruitment to localized plasma membrane regions (white arrow) upon localized activation of blue opsin using a confined 488 nm light pulse (blue box). Changing the direction of the 488 nm light results in EndoGi Tracker recruitment to the opposite side (grey arrow). Upon exposing the whole cell to light, the sensor is recruited to the plasma membrane globally (yellow arrows). The kymographic view shows the EndoGi Tracker recruitment dynamics on the plasma membrane across time. The single-cell line plot shows EndoGi Tracker dynamics in response to localized 488 nm light. (b) In HeLa cells expressing α2AR-CFP, EndoGi Tracker, CRY2-mCh-RGS4Δ, and CIBN-CAAX, localized blue light to one side of the cell results in the recruitment of RGS4Δ (220 s). Addition of norepinephrine to this cell results in EndoGi Tracker recruitment to the opposite side of the cell (220 s, white arrows). CRY2-mCh-RGS4Δ recovers back to the cytosol upon keeping the cell in the dark (1800 s), which resulted in the sensor being recruited globally (yellow arrows). Next, the direction of blue light was reversed, resulting in localized RGS4Δ recruitment. We observed the disappearance of the sensor from the RGS4Δ-recruited side of the cell and the increase of sensor accumulation on the opposite side (200 s, grey arrows). The kymographic view shows that EndoGi Tracker recruitment occurs on the opposite side of the cell upon RGS4Δ recruitment to one side of the cell. The single-cell line plot shows EndoGi Tracker dynamics in response to localized RGS4Δ-mediated Gαi-GTP hydrolysis. (c) A representative diagram depicting signaling repercussions due to localized RGS4Δ recruitment by blue light and simultaneous receptor activation (Created with BioRender.com). (d) NCI-H125 cells expressing melanopsin-A333T mutant and the EndoGq Tracker were exposed to localized blue light while imaging cells with 594 nm red light. The confined blue light induced localized recruitment of the sensor (white arrows). Upon changing the direction of the blue light, the sensor also showed recruitment to the opposite side (grey arrows). The EndoGq Tracker showed global recruitment to the plasma membrane upon exposing the whole cell to blue light (yellow arrows). The kymographic view shows the EndoGq Tracker recruitment to the plasma membrane across time. The single-cell line plot shows EndoGq Tracker dynamics in response to localized melanopsin-A333T activation. Although data are shown from only one cell, experiments were conducted in multiple cells to test reproducibility.

Next, we further examined EndoGi Tracker’s detectability in subcellular GαiGTP dynamics in response to a GTPase specific to GαiGTP. Regulator of G protein signaling 4 domain (RGS4Δ), which accelerates GTP hydrolysis on GαiGTP, inducing heterotrimer reformation. Here, we utilized an optogenetic version of RGS4Δ to optically control its GTPase activity.^83^ RGS4Δ is tethered to an optically sensitive cryptochrome 2 (CRY2) and mCherry (for visualization) (CRY2-mCh-RGS4Δ). CRY2 dimerizes with its plasma membrane-targeted binding partner, a truncated version of cryptochrome-interacting basic-helix-loop-helix (CIBN-CAAX), upon blue light stimulation.^83^ Here, we expressed α2AR-CFP, CRY2-mCh-RGS4Δ, CIBN-CAAX, and EndoGi Tracker in HeLa cells. Before blue light or ligand addition, the EndoGi Tracker and CRY2-mCh-RGS4Δ were cytosolic (Fig. 6b, 0 s). While imaging cells with mCherry (to visualize CRY2-mCh-RGS4Δ) and Venus (to visualize the EndoGi Tracker), we exposed a localized region of the plasma membrane to 445 nm blue light (Fig. 6b, blue box). The localized blue light recruited CRY2-mCh-RGS4Δ to one side of the cell (Fig. 6b, 220 s). We then activated α2AR globally by adding 100 µM norepinephrine. The EndoGi Tracker accumulated at the opposite side from the CRY2-mCh-RGS4Δ localization (Fig. 6b, 220 s, and white arrows and plot: Side A). This data shows that, due to the localized GTPase activity of RGS4Δ, the cells could only show GαiGTP generation at the opposite side. Termination of blue light resulted in CRY2-mCh-RGS4Δ recovering to the cytosol (Fig. 6b, 1800 s), while the sensor exhibited global plasma membrane recruitment (Fig. 6B, 1800 s, and yellow arrows). Next, we exposed the opposite side of the cell to blue light. The resultant CRY2-mCh-RGS4Δ recruitment (Fig. 6b, 2000 s) also induced a significant loss of Venus fluorescence locally, indicating a localized GαiGTP hydrolysis due to RGS4Δ activity and EndoGi Tracker remaining on the opposite side (Fig. 6b, 2000 s, grey arrows and plot: Side B). Figure 6c demonstrates a diagram of EndoGi Tracker and cellular G protein dynamics upon CRY2-mCh-RGS4Δ localization with blue light to one side of the cell. Collectively, this data clearly demonstrates the feasibility of using the EndoGi Tracker to detect dynamics of subcellular GPCR-G protein activities.

Next, we examined the subcellular GαqGTP detection ability of the Venus-tagged EndoGq Tracker. To our knowledge, there are no Gq-coupled opsins that can resist activation by yellow light. Therefore, we replaced the fluorescence tag of the sensor with mCherry and created MAS-mCh-GRK2-RH, allowing us to image the EndoGq Tracker-mCh with red light (594 nm excitation) while photoactivating the opsin with blue light. For this experiment, we employed the recently developed Gq-coupled, spectrally blue-shifted melanopsin mutant, A333T, as WT melanopsin is activated by 594 nm light, whereas the A333T mutant is not.^80^ Here, we expressed melanopsin-A333T (untagged) and EndoGq Tracker-mCh in NCI-H125 cells. At the same time, imaging cells with 594 nm red light, a localized region of the cell was exposed to blue light (blue box). Here, the localized melanopsin-A333T activation induced asymmetric GαqGTP generation on the blue-light-exposed side of the cell, resulting in localized recruitment of the EndoGq Tracker-mCh (Fig. 6d, 52 s, white arrow, and line plot: Side A). Upon switching the direction of the blue light to the opposite side, the sensor was also recruited to the opposite side (Fig. 6d, 1036 s, grey arrow, and line plot: Side B). Upon exposure of the whole cell to blue light, the EndoGq Tracker was globally recruited to the plasma membrane (Fig. 6d, 1980s, and yellow arrows, and plot). This data indicates the sensor’s ability to detect subcellular GαqGTP generation.

### Conclusions

Here, we describe the engineering and use of EndoGi and EndoGq Trackers, which detect endogenous, location-specific, dynamic activities of GαiGTP and GαqGTP. Compared to the currently used biosensors for G proteins, which require overexpression of the desired G protein, tagging large fluorescent or luminescent proteins to the protein of interest, or lacking subcellular resolution, our endogenous GαGTP sensors will be advantageous for dissecting G protein signaling mechanisms without perturbing the genetic makeup of the associated pathway. Moreover, the sensor design methods used in this study to selectively target probes to the plasma membrane using lipid motifs and lipid-interacting chaperones are innovative and can be adapted and optimized to engineer new biosensors.

Demonstrating the utility of our sensors, here we show that Gi-GPCR activation-induced heterotrimer activation is Gγ subtype-dependent. We have previously shown that Gβγ regulates effector activity in a Gγ-subtype-dependent manner, including PI3K-PIP3 generation,^31^ PLCβ-PIP2 hydrolysis,^44^ and cell migration.^84^ However, no studies have shown that the Gγ subtype on the Gβγ dimer modulates the entire heterotrimer activity. Gγ shows cell-tissue-subtype specific expression levels.^31,43^ Our data generated using the new sensors show that low-membrane affinity Gγ-subtype (e.g., Gγ9) expression in cells results in a faster and higher turnover of heterotrimer activation compared to low-membrane affinity Gγ (e.g., Gγ3) expressing cells. Since GPCR-G protein pathways are among the major drug targets in pharmacology,^4,73,85-87^ our findings could help understand why a specific GPCR-targeting drug may take longer to respond in a particular tissue than in another. Based on the above findings, we propose the hypothesis that Gγ composition of a cell is a predictor of GPCR-G protein signaling efficacy.

The Gq pathway is implicated in many diseases, including cancer and cardiovascular diseases.^30,38^ Therefore, understanding the molecular-level regulation of Gq pathway signaling is not only fundamental in understanding pathophysiology but also in therapy development. The newly engineered GαqGTP sensor enables the spatially resolved measurement of Gq pathway-induced heterotrimer activity in living cells. Our experimental data from living cells clearly show robust GαqGTP generation and subsequent adaptation upon Gq-GPCR activation. Given that GNAQ is a cancer driver and our data indicate rapidly adapting GαqGTP levels, we propose that cells likely employ a novel regulatory mechanism to protect from excessive Gαq signaling. Collectively, our sensors will help provide a dynamic molecular picture of endogenous GαGTP signaling dynamics at subcellular resolution and may reveal novel regulatory mechanisms masked under overexpression conditions, which are crucial to understanding pathological processes.

## Methods

### Reagents

The reagents used were as follows: Norepinephrine (NE) (Sigma Aldrich), Bombesin (Tocris Bioscience), 11-cis-retinal (National Eye Institute), Carbachol (Cayman Chemical), WIN-55,212-2 (Cayman Chemical), AM251 (Cayman Chemical), Fentanyl (Covetrus), and Naloxone (Covetrus). Stock solutions of compounds were prepared according to manufacturers’ recommendations and diluted to working concentrations in regular cell culture medium before adding to cells. Working concentrations of all reagents are provided in the specific experimental results.

### DNA constructs

The DNA constructs used were as follows: GRPR, α2AR-CFP, Blue opsin-mTurquoise, αq–CFP, mCh-Gγ9, and mCh-Gγ3 have been described previously.^29,44,77,88,89^ Venus-mini-G constructs were kindly provided by Dr. Nevin Lambert’s laboratory, Augusta University, Augusta, GA. MOR-mCh plasmid DNA was gifted by Dr. Zhou-Feng Chen at Washington University Pain Center. Dr. Susruta Majumdar from Washington University, Saint Louis, kindly provided us with the plasmid encoding CB1R. All EndoG(i/q) Tracker constructs were cloned by Gibson assembly cloning (NEB) using the BERKY DNA constructs for Gi and Gq, kindly provided by Mikel Garcia-Marcos, Boston University, MA. Cloned cDNA constructs were confirmed by Sanger sequencing (Genewiz).

### Cell culture and transfections

Cell lines used were as follows: HeLa and NCI-H125 cells were purchased from the American Type Culture Collection (ATCC). NCI-H125 cells were cultured in Roswell Park Memorial Institute (RPMI) 1640 medium (Corning, Manassas, VA) supplemented with 10% heat-inactivated dialyzed fetal bovine serum (DFBS, Atlanta Biologicals, GA) and 1% penicillin-streptomycin (PS, 10,000 U/ml stock) and grown at 37 °C with 5% CO_2_. Cells were cultured 60 mm or 100 mm cell culture dishes (Celltreat). Two days prior to imaging, 8 × 10^4^ cells were seeded on a 14 mm glass-bottomed well with a #1.5 glass coverslip in a 29 mm cell culture dish. The following day, cells were transfected with appropriate DNA combinations using the transfection reagent, Lipofectamine 2000 (Invitrogen) according to the manufacturer’s protocol and then incubated in a 37 °C, 5% CO2 incubator. Cells were imaged after 16 h post-transfection.

### Live cell imaging, image analysis, and data processing

The methods, protocols, and parameters for live-cell imaging are adapted from previously published work.^90-92^ Briefly, live-cell imaging experiments were performed using a spinning disk (Yokogawa CSU-X1, 5000 rpm) XD confocal TIRF imaging system composed of a Nikon Ti-R/B inverted microscope with a 60X, 1.4 NA oil objective and an iXon ULTRA 897BV back-illuminated deep-cooled EMCCD camera. Photoactivation and spatio-temporally controlled light exposure on cells in regions of interest (ROI) were performed using a laser combiner with 40-100 mW solid-state lasers (445, 488, 515, and 594 nm) equipped with an Andor® FRAP-PA unit (fluorescence recovery after photobleaching and photoactivation), controlled by Andor iQ 3.1 software (Andor Technologies, Belfast, United Kingdom). Fluorescent sensors, mCh-γ3, mCh-γ9, mCh-tagged EndoGi/Gq tracker constructs, mCh-PH, mCh-GRPR, and CRY2-mCh-RGS4Δ were imaged using 594 nm excitation−624 nm emission settings; all Venus-tagged EndoG(i/q) Tracker constructs and miniG constructs were imaged using 515 nm excitation and 542 nm emission; α2AR-CFP, Gαq-CFP, and Blue opsin-mTq were imaged using 445 nm excitation and 478 nm emission. Before experiments, we selected cells expressing α2AR or blue opsin GPCR by imaging the receptors with the 445 nm laser (before adding the ligand or retinal). Additional adjustments of laser power with 0.1%-1% transmittance were achieved using Acousto-optic tunable filters (AOTF). Data acquisition was performed using Andor iQ 3.1 software, as previously described.^90^ Time-lapse images were analyzed using Fiji (ImageJ, NIH) to quantify plasma membrane fluorescence over time. A custom in-house Macro script was used to automate membrane fluorescence quantification (Macro code is provided in the Supplementary Information). The script generates regions of interest (ROIs) on the plasma membrane and extracts mean fluorescence intensity values across all time points. Further data processing, baseline normalization, and statistical analysis were performed as published in previous studies.^31,80,90^

### Computational modeling and docking

AlphaFold2^62^ was used to predict homology models of all sensor structures. Coverage scores were reported as the Predicted Local Distance Test (pLDDT) and predicted template modeling score (pTM), which estimated the accuracy of the local structure of each amino acid residue and the accuracy of the global structure, respectively. The models with the highest coverage scores (Rank No. 1) were selected for further in-silico analysis. PyMOL3.1+ was used to align each domain of the modeled AlphaFold2 structures to existing PDB structures (5TB5, 5L7K, 7OK6, 5IBO, 3AKO, and 2BCJ) to ensure that each domain is conserved and folded/oriented properly.

The Schrödinger Maestro (Schrödinger Release 2025-4: Maestro, Schrödinger, LLC, New York, NY, 2025) was used to perform docking studies for the predicted homology models. Each model structure was prepared for docking using the Protein Preparation wizard in the Schrödinger module to optimize hydrogen bonding networks and remove steric clashes. The structure of the myristoyl group was prepared using the Ligprep wizard. Next, the myristoyl group was docked to the protein models using the GLIDE tool in Schrödinger. Hydrophobic cavities corresponding to the predicted myristoyl binding sites were identified via SiteMap analysis. Docked poses were evaluated based on Docking Score and Glide Emodel values represented in kcal/mol. The structure with the highest negative docking score and the Glide Emodel value (Site No. 1) was selected as the most favorable myristoyl group binding site in the analyzed proteins.^64,65^

### Statistical Data Analysis

All experiments were repeated multiple times (>3) to test the reproducibility of the results. Statistical analysis and data plotting were performed using OriginPro (OriginLab®). Results were analyzed from multiple cells and represented as mean ± SD. The exact number of cells used in the analysis is given in the figure legends. The sensor recruitment and recovery rates were calculated using the Nonlinear Curve Fitting tool (NLFit) in OriginPro. Each plot was fitted to the DoseResp (Dose-Response) function under the Pharmacology category in OriginPro. The mean values of hill slopes (P) obtained for each nonlinear curve fitting are presented as mean rates. One-way ANOVA tests were performed in OriginPro to determine statistical significance between two or more populations of signaling responses. Tukey’s mean-comparison test was performed at the p < 0.05 significance level for the one-way ANOVA. All statistical test results are given in the Supplementary Information.

## Author contributions

D.W conceived and designed the sensors, and performed most live cell imaging experiments, and the data analysis. D. W. And S.P. performed plasmid cloning. D.W. developed the custom Macro code for automated plasma membrane quantification in Fiji. S.P. conducted opioid receptor experiments, subcellular experiments, and tested the reproducibility of the experimental data. A.K. and D.W conceptualized the project and wrote the manuscript.

## Acknowledgements

We acknowledge Dr. Mikel Garcia-Marcos, Dr. Nevin Lambert, Dr. Zhou-Feng Chen, and Dr. Susruta Majumdar for providing us with plasmid DNA constructs essential for this study. We thank the National Eye Institute for providing 11-cis-retinal. We are grateful to the Department of Comparative Medicine of Saint Louis University for providing the controlled substances for opioid receptor activation and the necessary licensing/ certifications. We are also grateful to the Institute for Drug and Biotherapeutic Innovation of Saint Louis University (SLU-IDBI) for providing liscence to use the Schrödinger suite and PyMOL3.1+ software for the in silico analysis. BioRender for providing the platform to create some of the figures.BioRender for providing the platform to create some of the figures.

## Code availability

The macro code used in this study for image analysis is provided in the Supplementary Information.

