## Supplementary Information for "Novel GαGTP Sensors Reveal Endogenous and Subcellular G Protein Signaling Dynamics"

#### Figure S1

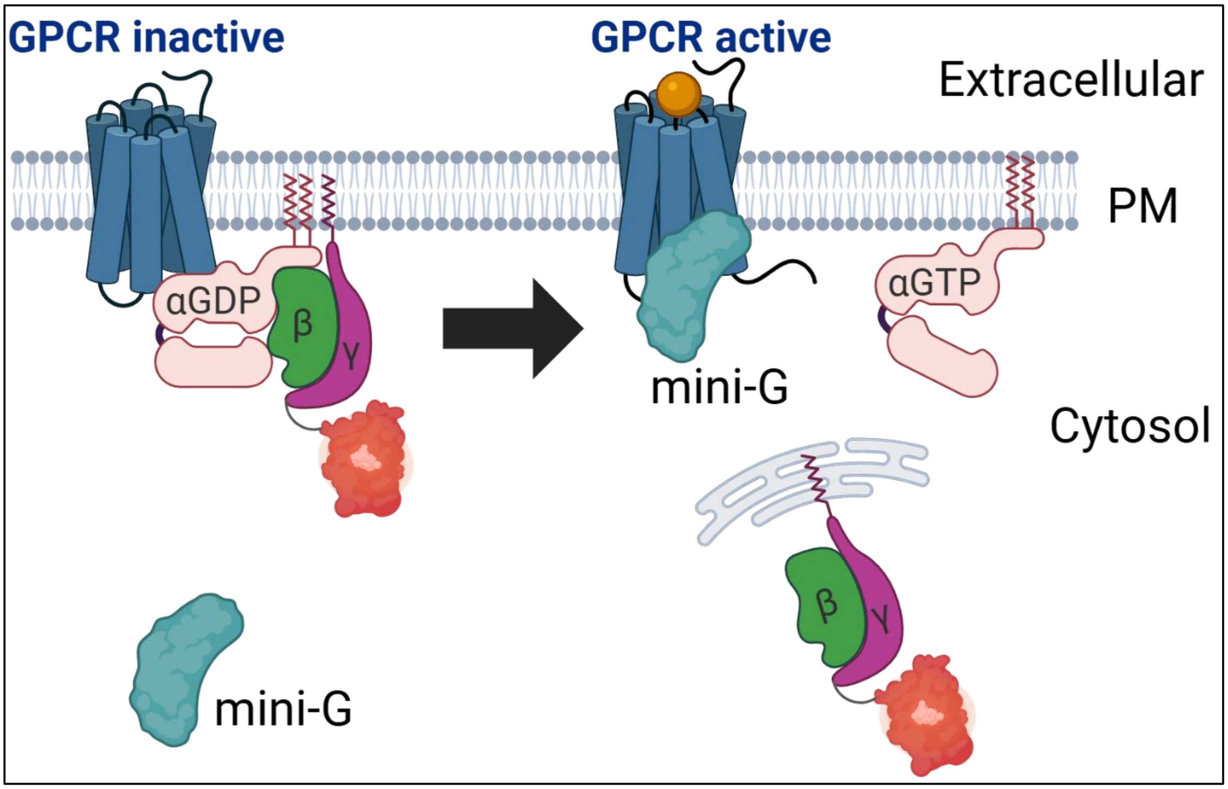


**Figure S1: Graphic representation of heterotrimer dissociation upon GPCR activation and assays used to probe active state GPCRs.** Cytosolic Mini-G proteins are recruited to the GPCR upon GPCR activation. Fluorescently-tagged Gβγ translocation to internal membranes upon heterotrimer dissociation indicates GPCR- G protein activity. Created with Biorender.com.

#### Figure S2

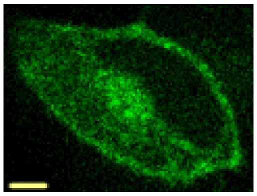

MAS-KB-1753-
PDEδ6-Venus

**Figure S2: MAS-KB-1753-PDEδ6-Venus shows plasma membrane localization.** HeLa cells expressing MAS-KB-1753-PDEδ6-Venus shows prominent plasma membrane localization indicating very low or no chaperoning of the myristoyl anchor by PDEδ6. The scale bar = 5 μm

#### Figure S3

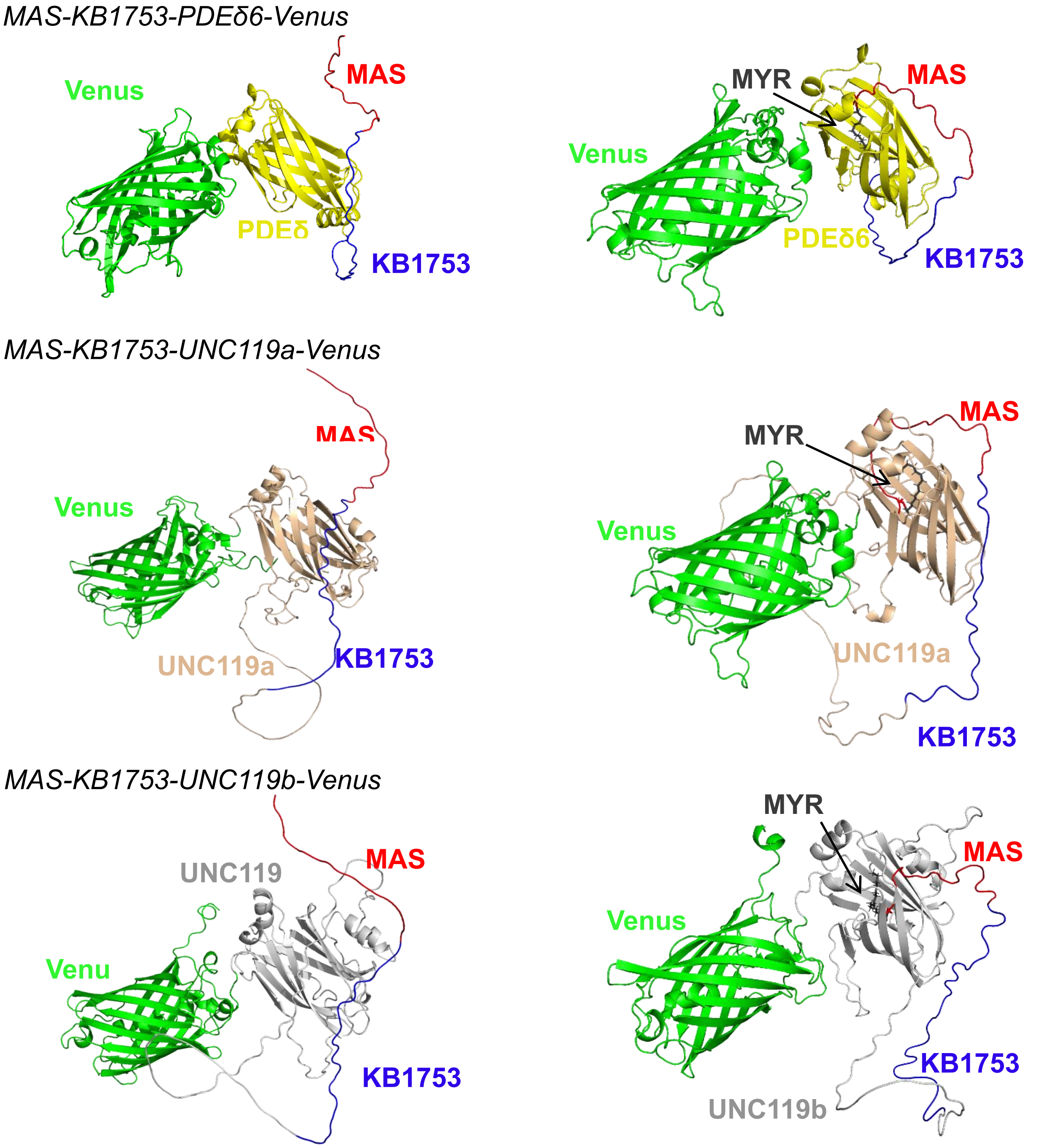


**Figure S3: *In-silico* analysis of interactions between myristoyl group and chaperone in failed sensors.** Homology models MAS-KB1753-PDEδ6-Venus, MAS-KB1753-UNC119a-Venus, and MAS-KB1753-UNC119b-Venus obtained from AlphaFold2 modeling software. The homology models with highest coverage scores are shown. Docking hits from the Schrödinger software with the best poses and the most favorable docking scores are shown here. The myristoyl group is docked into the hydrophobic pockets of chaperones. Different regions of the protein are depicted in separate colors; MAS: red; KB1753: dark blue; Venus: green; myristoyl group: black; PDEδ6: yellow; UNC119a: wheat; UNC119b: grey.

#### Figure S4

a

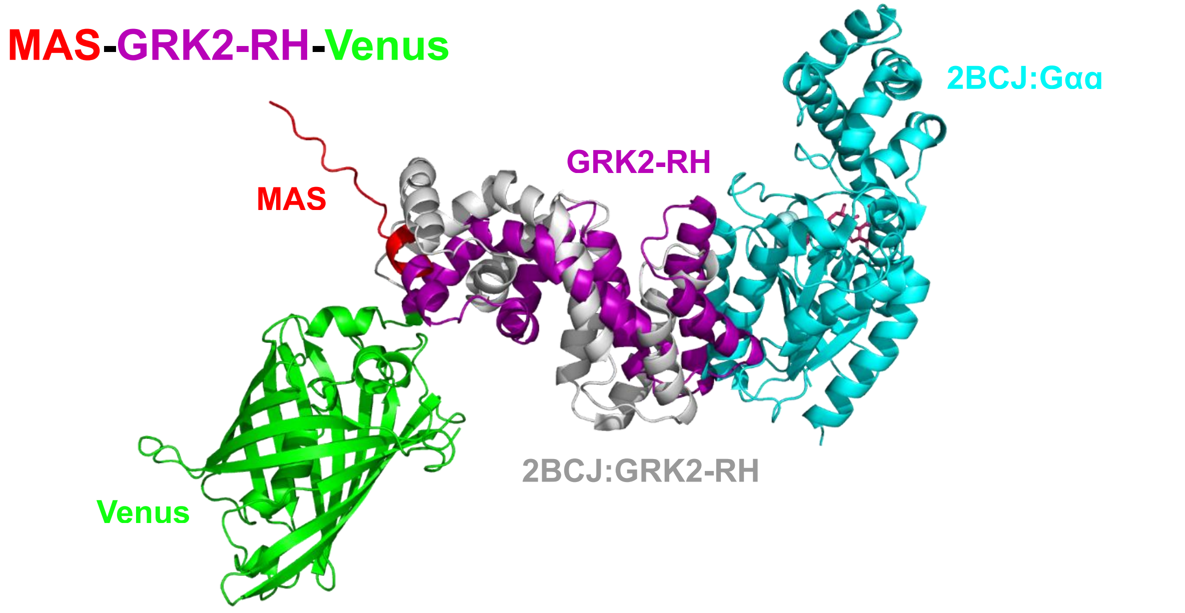


b

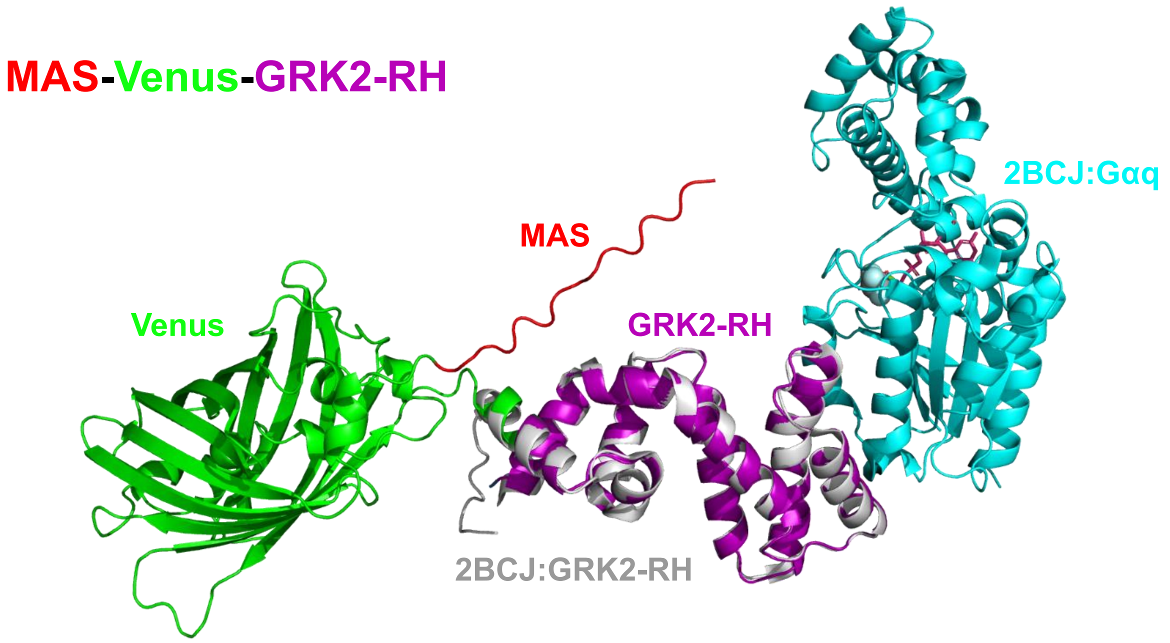


**Figure S4: *In-silico* analysis of MAS-GRK2-RH--Venus and MAS-Venus-GRK2-RH orientations. (a)** MAS-GRK2-RH-Venus is aligned with GRK2 bound Gαq (PDB:2BCJ) using Pymol software. GRK2-RH domain in the modeled sensor is displaced with GRK2-bound to Gαq indicating folding/ orientation issues of the protein. **(b)** MAS-Venus GRK2-RH aligned with GRK2 bound Gαq (PDB:2BCJ) using Pymol software. The alignment shows near-perfect overlay between the GRK2-RH domains in the two protein structures. This suggests efficient orientation/ folding for sensor activity.

### Figure S5

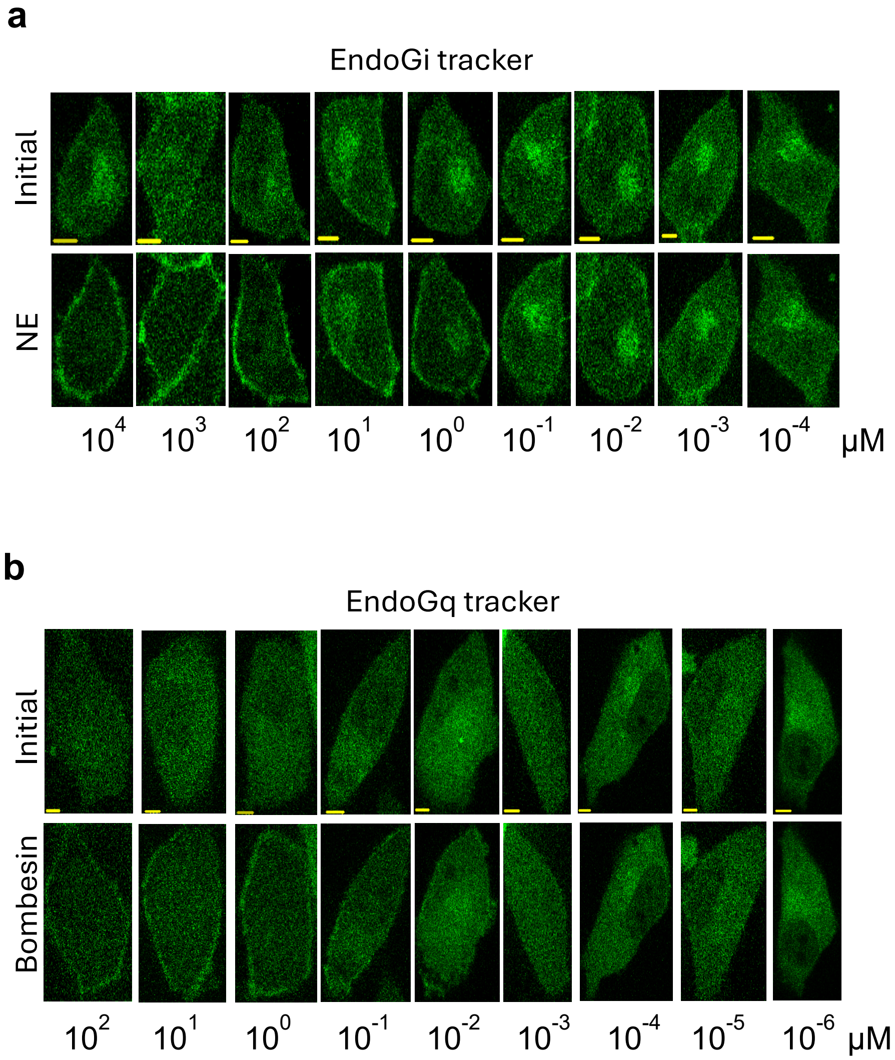


**Figure S5: EndoG(i/q) Trackers show dose dependent activity curves. (a)** Cell images show average response of EndoGi Tracker to α2AR activation to norepinephrine concentration gradient of 0.0001-10000 µM. **(b)** H125 cell images showing the average response of EndoGq Tracker upon activation of GRPR via a Bombesin concentration gradient (0.001 nM-100 uM). The scale bar = 5 µm

### Figure S6

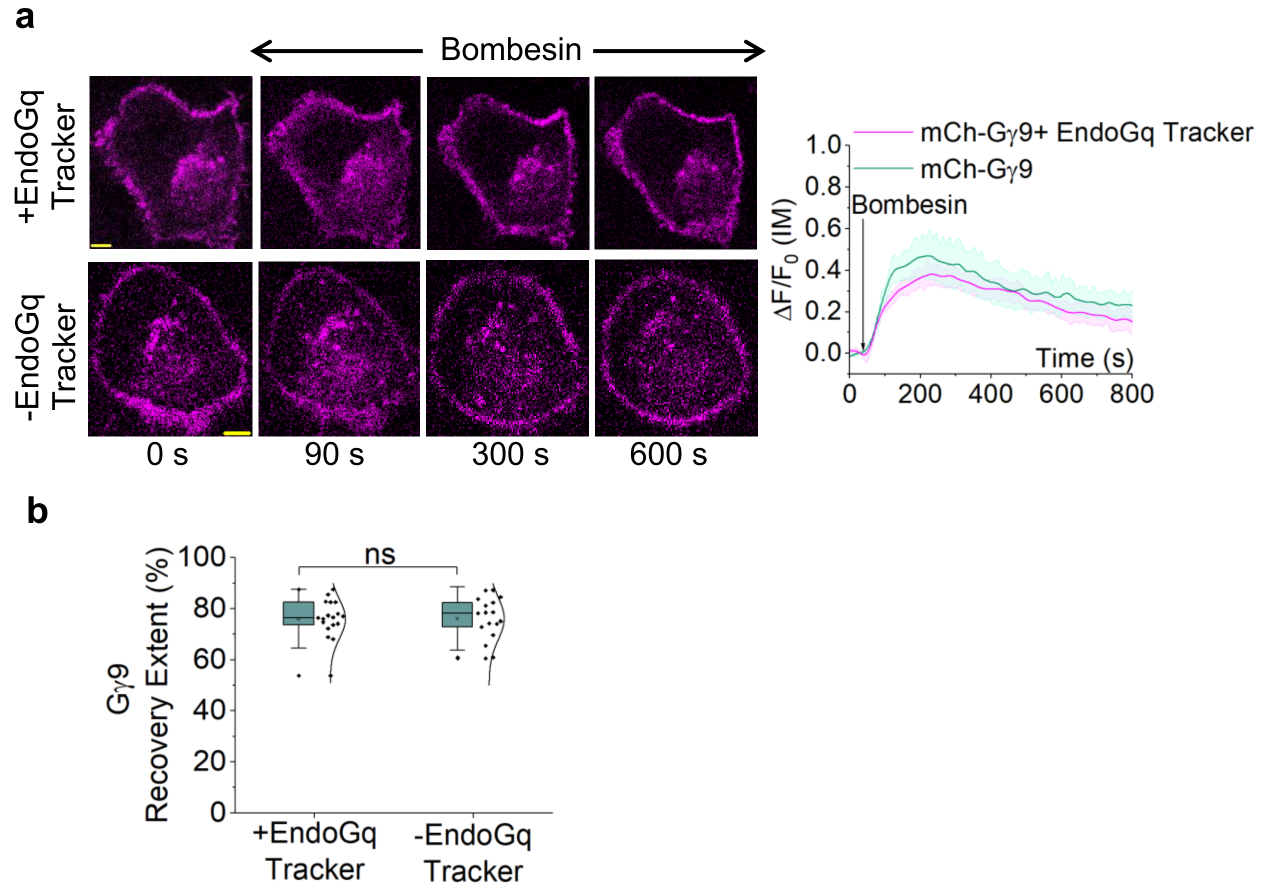


**Figure S6: EndoGq Tracker depicts molecular details underlying Gq signaling adaptation. (a)** NCI-H125 cells show robust mCh-Gγ9 translocation and subsequent recovery independent of EndoGq Tracker expression. **(b)** The Gγ9 recovery extents are not different between native NCI-H125 ($n = 17$) and cells expressing EndoGq Tracker ($n = 18$). IM: Internal Membrane; The scale bar = 5 µm. Statistical comparisons were performed using One-way-ANOVA; $p > 0.05$, (ns: population means are not significantly different at 95% confidence level).

### Figure S7

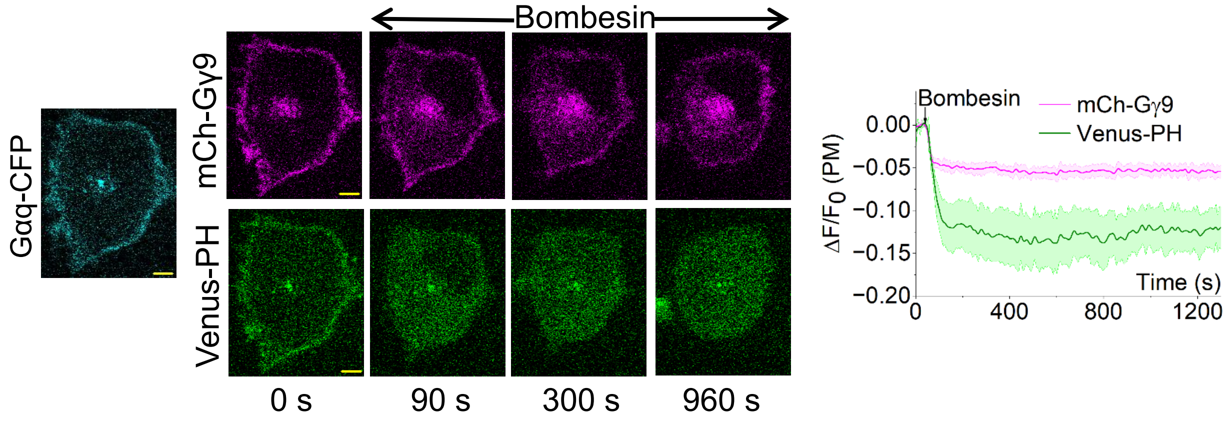


**Figure S7: NCI-H125 cells expressing Gαq-CFP shows non-adapting G protein activation and downstream signaling.** NCI-H125 cells expressing GRPR, Gαq-CFP, mCh-Gγ9, and Venus-PH shows robust Gγ9 translocation and PIP2 hydrolysis upon GRPR activation and stays translocated. The hydrolyzed PIP2 and translocated Gγ9 did not recover back to the plasma membrane ($n = 10$). The scale bar = 5 μm

#### Figure S8

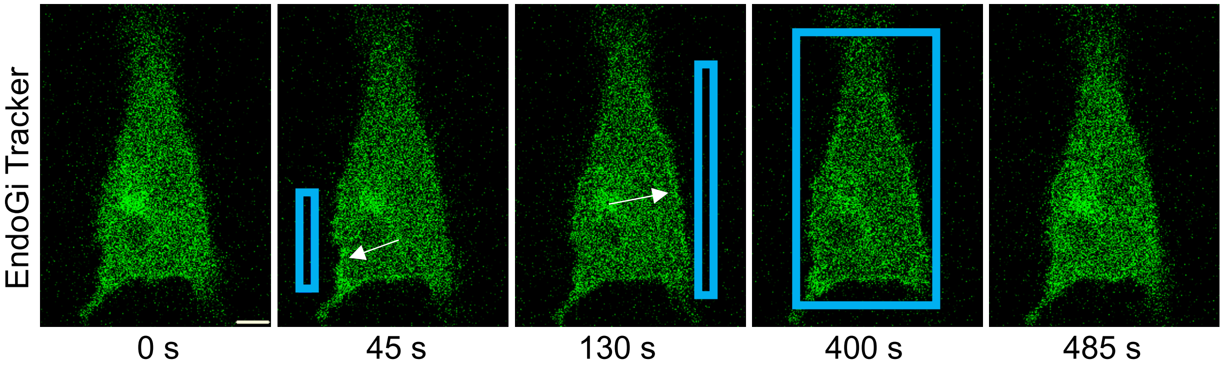


**Figure S8: The optogenetic utility of the EndoGi Tracker with bistable opsins.** HeLa cells expressing Lamprey parapinopsin and EndoGi Tracker exhibit localized sensor recruitment in response to photoactivation of the receptor by blue light. The blue box indicates the direction of blue light. The scale bar = 5 μm

#### Figure S9

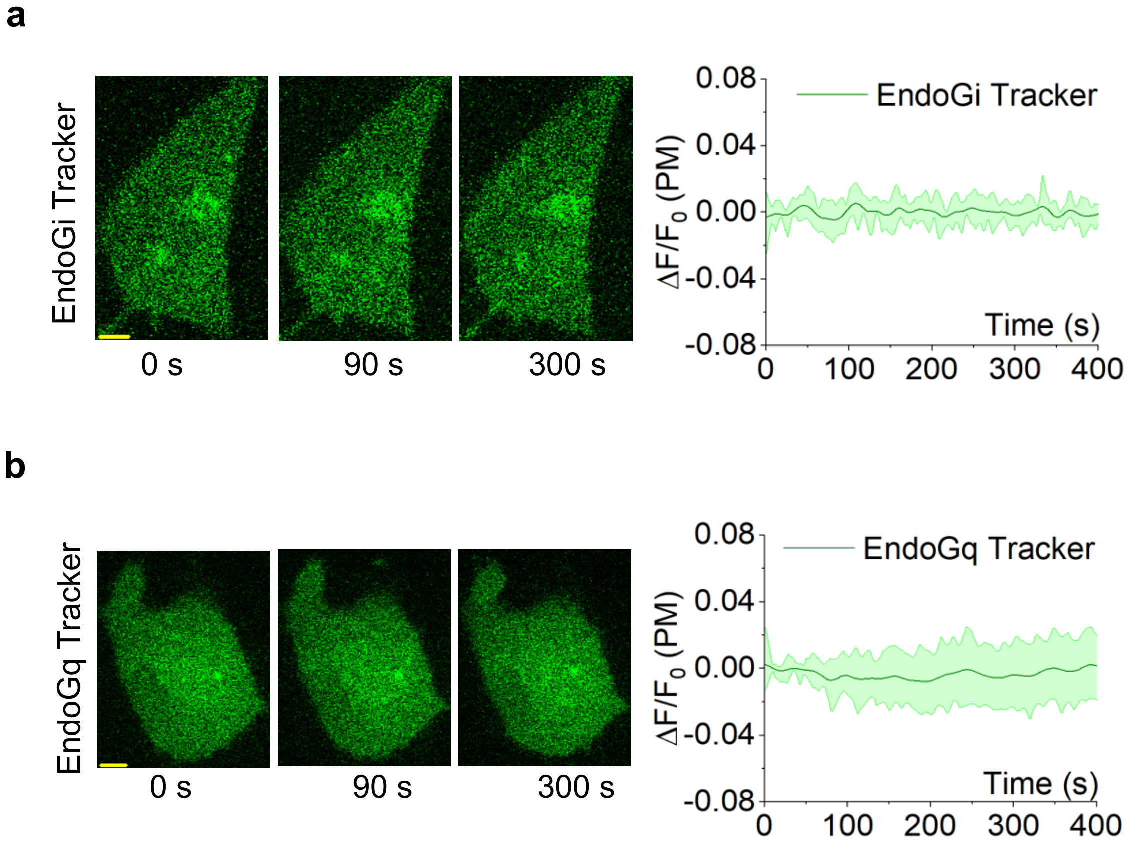


**Figure S9: Negative control experiments show no recruitment of EndoG(i/q) Trackers in the absence of receptor activation. (a)** EndoGi Tracker shows no plasma membrane recruitment without norepinephrine addition in HeLa cells expressing α2AR (n = 12). (b) No plasma membrane recruitment of EndoGq Tracker was observed without bombesin to activate GRPR in NCI-H125 cells (n = 13). The scale bar = 5 µm.

**Supplementary Table 1: Coverage scores for MAS-KB1753-PDEδ6-Venus homology models by AlphaFold2**

| Rank No. | Structure | PDB B factor (pLDDT) | pTM |
|---|---|---|---|
| 1 | 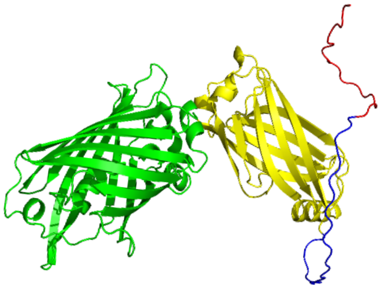 | 87.1 | 0.668 |
| 2 | 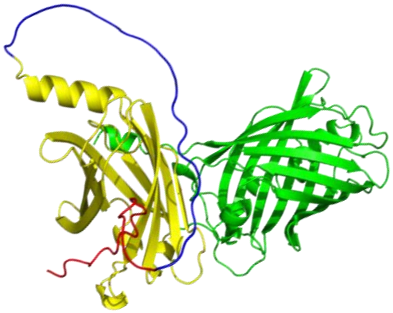 | 86.9 | 0.635 |
| 3 | 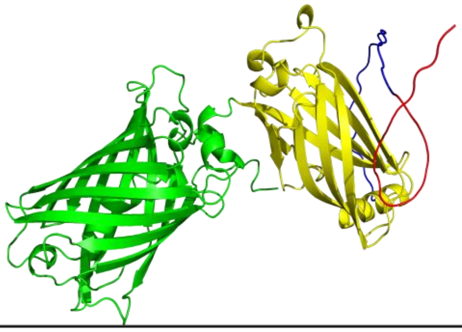 | 86.6 | 0.635 |
| 4 | 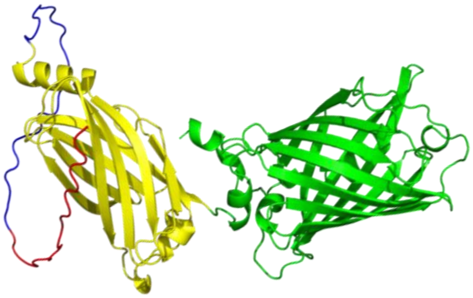 | 86.1 | 0.633 |
| 5 | 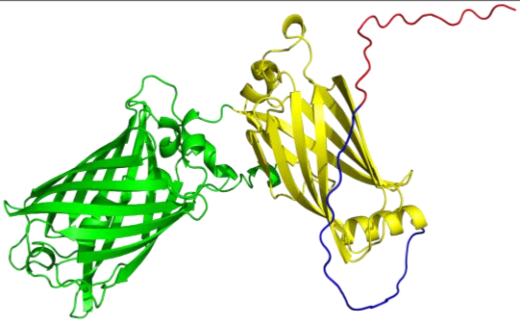 | 86.1 | 0.610 |

**Supplementary Table 2: Molecular Docking results for MAS-KB1753-PDEδ6-Venus by the Schrödinger Maestro software**

| Site No. | Docking score | Glide emodel kcal/mol | Structure |
|---|---|---|---|
| 1 | -0.778 | -25.392 | 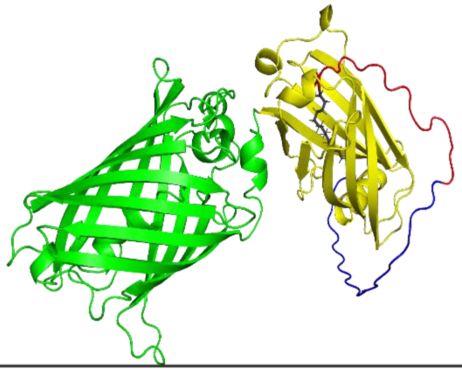 |
| 2 | 0.923 | -23.025 | 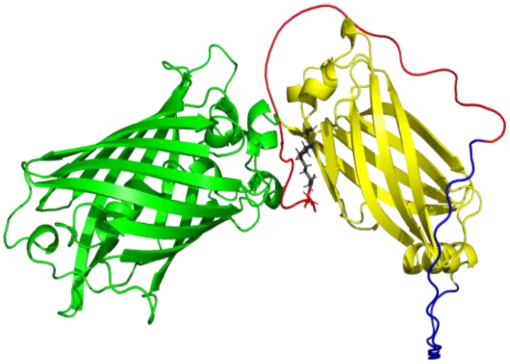 |
| 3 | 1.961 | -20.100 | 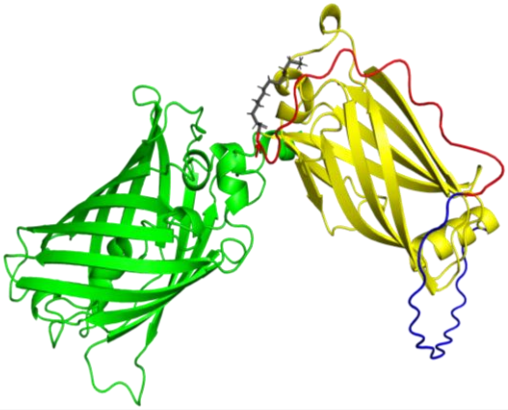 |
| 4 | 1.016 | -21.972 | 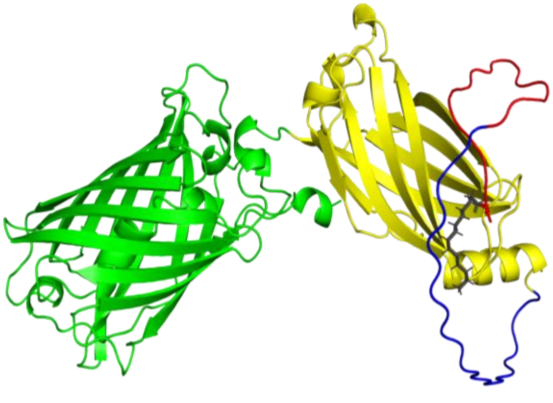 |

**Supplementary Table 3: Coverage scores for MAS-KB1753-UNC119a-Venus homology models by AlphaFold2**

| Rank No. | Structure | PDB B factor (pLDDT) | pTM |
|---|---|---|---|
| 1 | 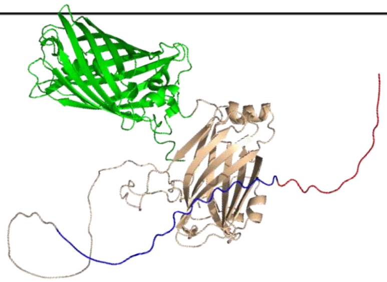 | 78.6 | 0.525 |
| 2 | 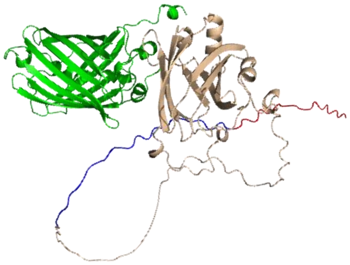 | 78.4 | 0.567 |
| 3 | 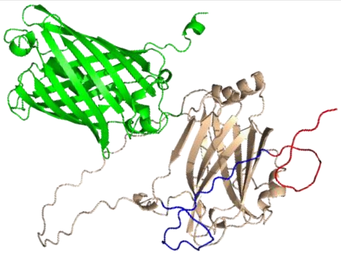 | 78.4 | 0.529 |
| 4 | 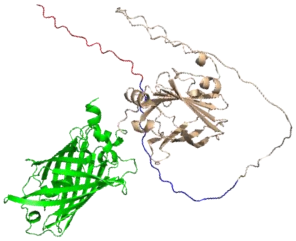 | 78.3 | 0.521 |
| 5 | 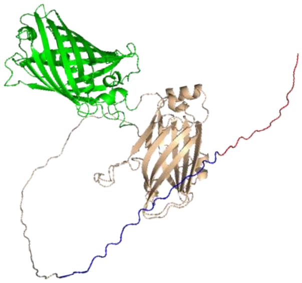 | 77.2 | 0.514 |

**Supplementary Table 4: Molecular Docking results for MAS-KB1753-UNC119a-Venus by the Schrödinger Maestro software**

| Site No. | Docking score | Glide emodel kcal/mol | Structure |
|---|---|---|---|
| 1 | -0.063 | -28.401 | 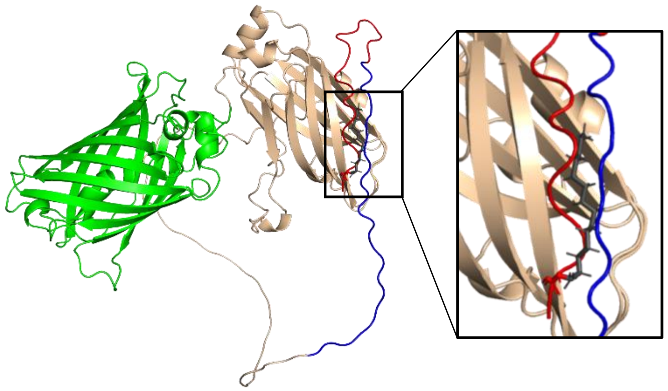 |
| 2 | -0.846 | -30.239 | 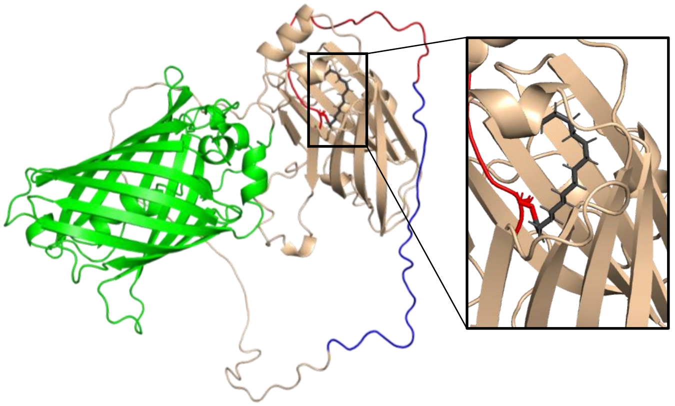 |
| 3 | -0.423 | -28.283 | 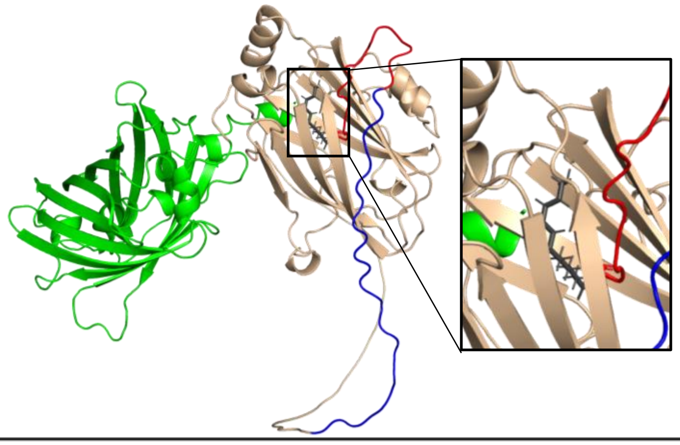 |

**Supplementary Table 5: Coverage scores for MAS-KB1753-UNC119b-Venus homology models by AlphaFold2**

| Rank No. | Structure | PDB B factor (pLDDT) | pTM |
|---|---|---|---|
| 1 | 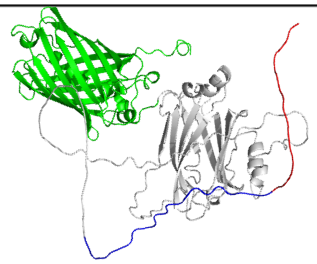 | 79.3 | 0.527 |
| 2 | 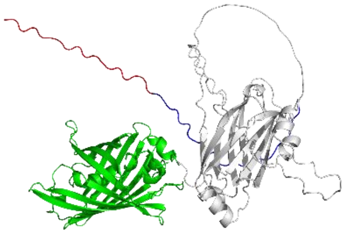 | 79.1 | 0.525 |
| 3 | 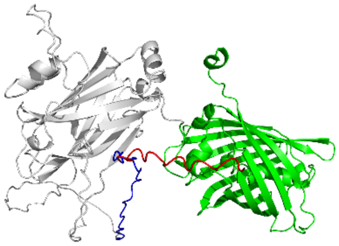 | 79.1 | 0.565 |
| 4 | 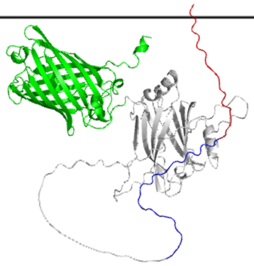 | 78.9 | 0.527 |
| 5 | 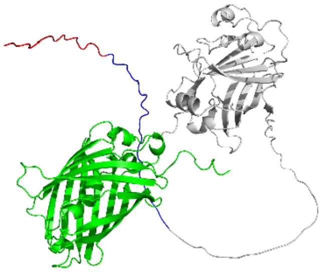 | 78.2 | 0.526 |

**Supplementary Table 6: Molecular Docking results for MAS-KB1753-UNC119b-Venus by the Schrödinger Maestro software**

| Site No. | Docking score | Glide emodel kcal/mol | Structure |
|---|---|---|---|
| 1 | -0.320 | -30.018 | 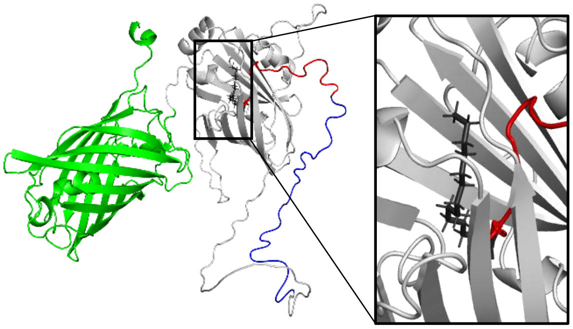 |
| 2 | 1.770 | -20.921 | 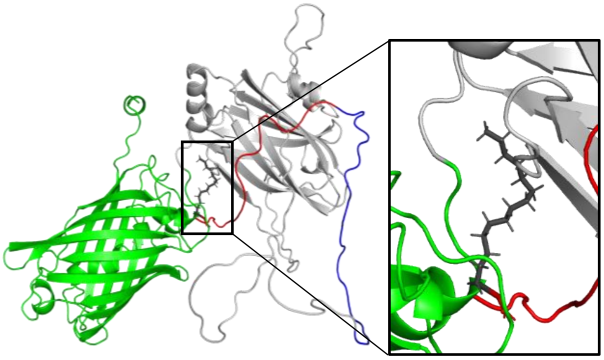 |
| 3 | 2.445 | -18.778 | 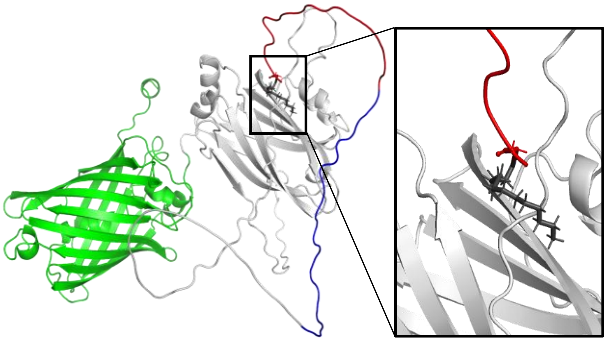 |
| 4 | 1.622 | -21.125 | 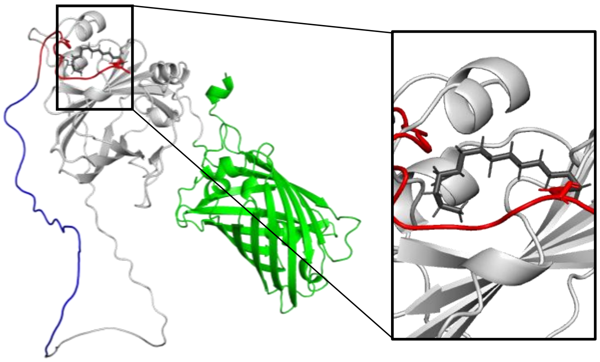 |

**Supplementary Table 7: Coverage scores for Docking results for MAS-KB1753-Nluc-Venus homology models by AlphaFold2**

| Rank No. | Structure | PDB B factor (pLDDT) | pTM |
|---|---|---|---|
| 1 | 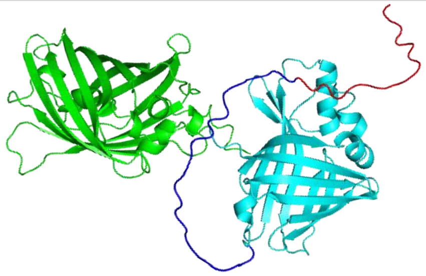 | 85.2 | 0.593 |
| 2 | 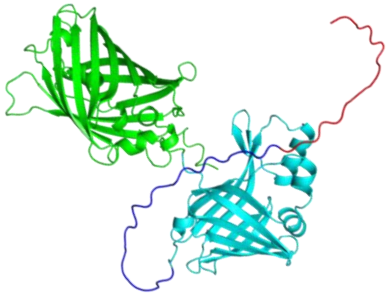 | 84.9 | 0.587 |
| 3 | 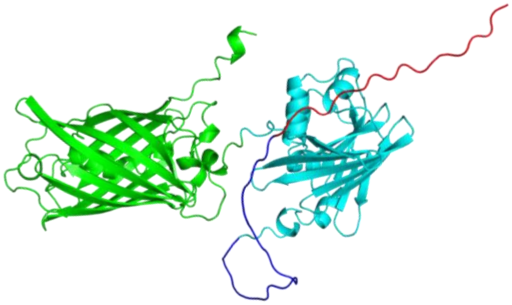 | 70.2 | 0.565 |
| 4 | 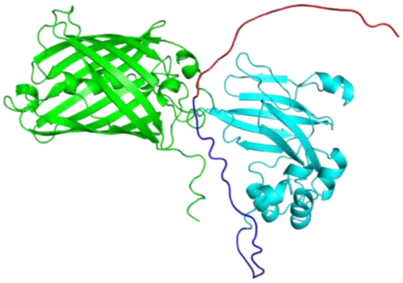 | 65.4 | 0.565 |
| 5 | 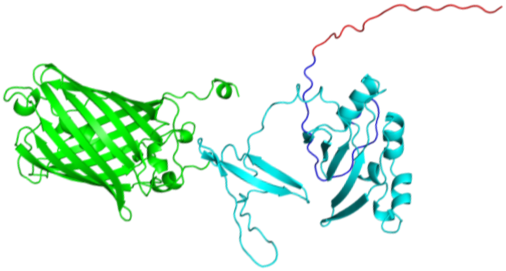 | 64.8 | 0.565 |

**Supplementary Table 8: Molecular Docking results for MAS-KB1753-Nluc-Venus by the Schrödinger Maestro software**

| Site No. | Docking score | Glide emodel kcal/mol | Structure |
|---|---|---|---|
| 1 | 1.460 | -19.538 | 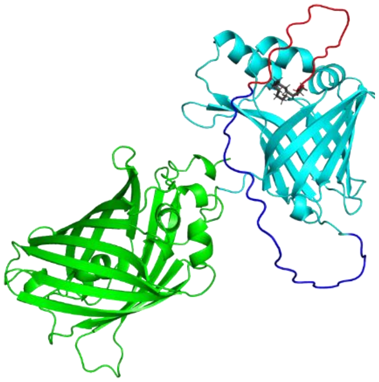 |
| 2 | 2.160 | -19.171 | 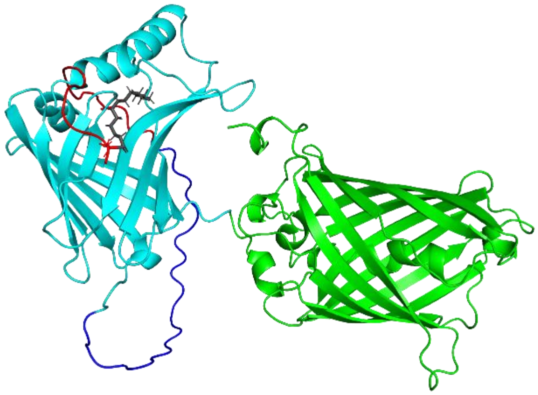 |
| 3 | 3.981 | 3.236 | 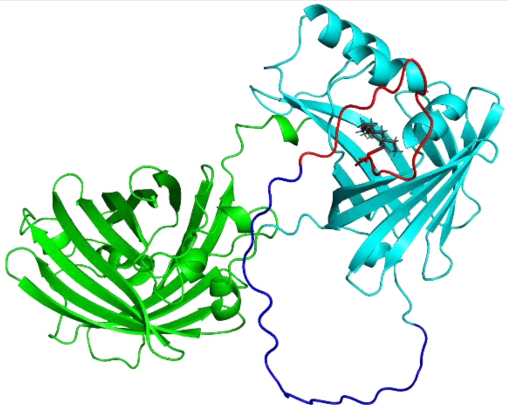 |

**Supplementary Table 9: Coverage scores for Docking results for MAS-Venus-GRK2-RH homology models by AlphaFold2**

| Rank No. | Structure | PDB B factor (pLDDT) | pTM |
|---|---|---|---|
| 1 | 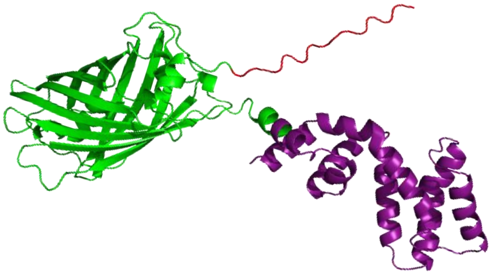 | 91.0 | 0.691 |
| 2 | 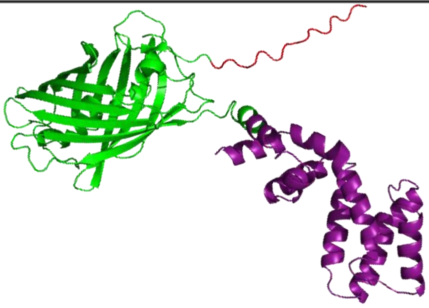 | 90.9 | 0.695 |
| 3 | 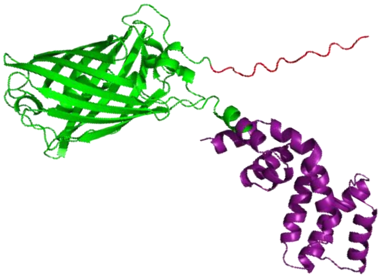 | 90.9 | 0.700 |
| 4 | 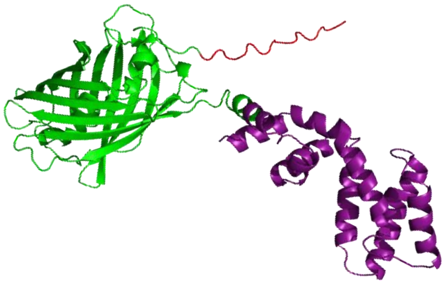 | 90.5 | 0.708 |
| 5 | 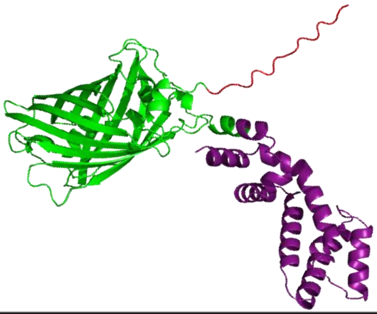 | 89.9 | 0.658 |

**Supplementary Table 10: Molecular Docking results for MAS-Venus-GRK2-RH by the Schrödinger Maestro software**

| Site No. | Docking score | Glide emodel kcal/mol | Structure |
|---|---|---|---|
| 1 | 2.505 | -12.771 | 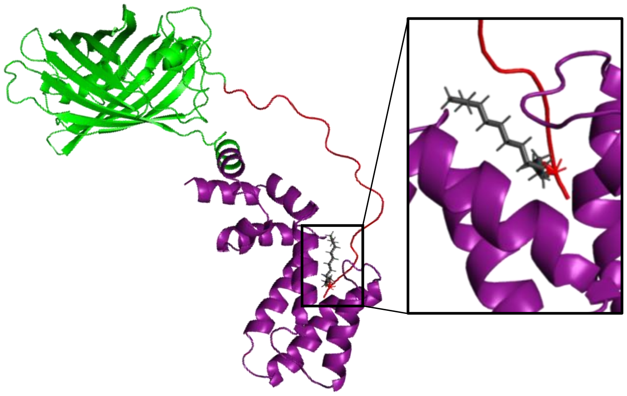 |

**Supplementary Table 11:** Descriptive Statistics for EndoGi Tracker recruitment extent with different chaperones

| | N Analysis | Mean | Standard Deviation | SE of Mean |
|---|---|---|---|---|
| UNC119a | 8 | 0.06798 | 0.03644 | 0.01288 |
| UNC119b | 9 | 0.02109 | 0.00926 | 0.00309 |
| NLuc | 9 | 0.13262 | 0.07645 | 0.02548 |

| | DF | Sum of Squares | Mean Square | F Value | Prob>F |
|---|---|---|---|---|---|
| Model | 2 | 0.05642 | 0.02821 | 11.4352 | 3.57E-04 |
| Error | 23 | 0.05673 | 0.00247 | | |
| Total | 25 | 0.11315 | | | |

At the 0.05 level, the population means are significantly different.

**Supplementary Table 12:** Descriptive Statistics for EndoGi Tracker recruitment and recovery with MOR

| | N Analysis | Mean | Standard Deviation | SE of Mean |
|---|---|---|---|---|
| Recruitment | 10 | 12.4349 | 3.22387 | 1.01948 |
| Recovery | 10 | 46.33779 | 12.36712 | 3.91083 |

| | DF | Sum of Squares | Mean Square | F Value | Prob>F |
|---|---|---|---|---|---|
| Model | 1 | 5747.03075 | 5747.03075 | 70.36935 | <0.0001 |
| Error | 18 | 1470.05125 | 81.66951 | | |
| Total | 19 | 7217.08199 | | | |

At the 0.05 level, the population means are significantly different.

**Supplementary Table 13:** Descriptive Statistics for EndoGi Tracker recruitment and recovery with CB1R

| | N Analysis | Mean | Standard Deviation | SE of Mean |
|---|---|---|---|---|
| Recruitment | 9 | 16.27003 | 2.58307 | 0.86102 |
| Recovery | 9 | 144.79442 | 36.49844 | 12.16615 |

| | DF | Sum of Squares | Mean Square | F Value | Prob>F |
|---|---|---|---|---|---|
| Model | 1 | 74333.3365 | 74333.3365 | 111.044 | <0.0001 |
| Error | 16 | 10710.46929 | 669.40433 | | |
| Total | 17 | 85043.80578 | | | |

At the 0.05 level, the population means are significantly different.

**Supplementary Table 14:** Descriptive Statistics for EndoGi Tracker recruitment rates with G$\gamma$3 and G$\gamma$9 co-expression

| | N Analysis | Mean | Standard Deviation | SE of Mean |
|---|---|---|---|---|
| With G$\gamma$3 | 11 | 0.046 | 0.03804 | 0.01147 |
| With G$\gamma$9 | 10 | 0.11352 | 0.09625 | 0.03044 |

| | DF | Sum of Squares | Mean Square | F Value | Prob>F |
|---|---|---|---|---|---|
| Model | 1 | 0.02387 | 0.02387 | 4.63551 | 0.04438 |
| Error | 19 | 0.09786 | 0.00515 | | |
| Total | 20 | 0.12173 | | | |

At the 0.05 level, the population means are significantly different.

**Supplementary Table 15:** Descriptive Statistics for EndoGi Tracker recruitment extent with Gγ3 and Gγ9 co-expression

| | N Analysis | Mean | Standard Deviation | SE of Mean |
|---|---|---|---|---|
| With Gγ3 | 11 | 0.03099 | 0.01225 | 0.00369 |
| With Gγ9 | 10 | 0.07092 | 0.03726 | 0.01178 |

| | DF | Sum of Squares | Mean Square | F Value | Prob>F |
|---|---|---|---|---|---|
| Model | 1 | 0.00835 | 0.00835 | 11.34076 | 0.00323 |
| Error | 19 | 0.014 | 7.36637E-4 | | |
| Total | 20 | 0.02235 | | | |

At the 0.05 level, the population means are significantly different.

**Supplementary Table 16:** Descriptive Statistics for EndoGi Tracker recovery rates with Gγ3 and Gγ9 co-expression

| | N Analysis | Mean | Standard Deviation | SE of Mean |
|---|---|---|---|---|
| With Gγ3 | 11 | 0.02034 | 0.01645 | 0.00496 |
| With Gγ9 | 10 | 0.03346 | 0.02079 | 0.00657 |

| | DF | Sum of Squares | Mean Square | F Value | Prob>F |
|---|---|---|---|---|---|
| Model | 1 | 9.01032E-4 | 9.01032E-4 | 2.59625 | 0.1236 |
| Error | 19 | 0.00659 | 3.47051E-4 | | |
| Total | 20 | 0.00749 | | | |

At the 0.05 level, the population means are not significantly different.

**Supplementary Table 17:** Descriptive Statistics for EndoGq Tracker recruitment extent with and without exogenous Gαq expression in HeLa cells

| | N Analysis | Mean | Standard Deviation | SE of Mean |
|---|---|---|---|---|
| With Gαq | 9 | 0.12737 | 0.05863 | 0.01954 |
| Without Gαq | 10 | 0.03228 | 0.01029 | 0.00325 |

| | DF | Sum of Squares | Mean Square | F Value | Prob>F |
|---|---|---|---|---|---|
| Model | 1 | 0.04283 | 0.04283 | 25.58755 | <0.0001 |
| Error | 17 | 0.02845 | 0.00167 | | |
| Total | 18 | 0.07128 | | | |

At the 0.05 level, the population means are significantly different.

**Supplementary Table 18:** Descriptive Statistics for EndoGq Tracker recruitment rates with and without exogenous Gαq expression in HeLa cells

| | N Analysis | Mean | Standard Deviation | SE of Mean |
|---|---|---|---|---|
| With Gαq | 9 | 0.09074 | 0.06593 | 0.02198 |
| Without Gαq | 10 | 0.01124 | 0.01016 | 0.00321 |

| | DF | Sum of Squares | Mean Square | F Value | Prob>F |
|---|---|---|---|---|---|
| Model | 1 | 0.02993 | 0.02993 | 14.25347 | 0.00151 |
| Error | 17 | 0.0357 | 0.0021 | | |
| Total | 18 | 0.06564 | | | |

At the 0.05 level, the population means are significantly different.

**Supplementary Table 19:** Descriptive Statistics for EndoGq Tracker's percent recovery with and without exogenous Gαq expression in HeLa cells

| | N Analysis | Mean | Standard Deviation | SE of Mean |
|---|---|---|---|---|
| With Gαq | 9 | 10.47162 | 12.15774 | 4.05258 |
| Without Gαq | 10 | 71.84977 | 20.73053 | 6.55557 |

| | DF | Sum of Squares | Mean Square | F Value | Prob>F |
|---|---|---|---|---|---|
| Model | 1 | 17844.99897 | 17844.99897 | 60.06895 | <0.0001 |
| Error | 17 | 5050.27969 | 297.07528 | | |
| Total | 18 | 22895.27866 | | | |

At the 0.05 level, the population means are significantly different.

**Supplementary Table 20:** Descriptive Statistics for EndoGq Tracker recruitment extent in NCI-H125 vs HeLa cells without exogenous Gαq expression

| | N Analysis | Mean | Standard Deviation | SE of Mean |
|---|---|---|---|---|
| NCI-H125 | 9 | 0.04642 | 0.02393 | 0.00798 |
| HeLa | 10 | 0.03228 | 0.01029 | 0.00325 |

| | DF | Sum of Squares | Mean Square | F Value | Prob>F |
|---|---|---|---|---|---|
| Model | 1 | 9.47201E-4 | 9.47201E-4 | 2.90943 | 0.10627 |
| Error | 17 | 0.00553 | 3.25563E-4 | | |
| Total | 18 | 0.00648 | | | |

At the 0.05 level, the population means are significantly different.

**Supplementary Table 21:** Descriptive Statistics for EndoGq Tracker recruitment rates in NCI-H125 vs HeLa cells without exogenous Gαq expression

| | N Analysis | Mean | Standard Deviation | SE of Mean |
|---|---|---|---|---|
| NCI-H125 | 9 | 0.14256 | 0.07114 | 0.02371 |
| HeLa | 10 | 0.01124 | 0.01016 | 0.00321 |

| | DF | Sum of Squares | Mean Square | F Value | Prob>F |
|---|---|---|---|---|---|
| Model | 1 | 0.08169 | 0.08169 | 33.52889 | <0.0001 |
| Error | 17 | 0.04142 | 0.00244 | | |
| Total | 18 | 0.1231 | | | |

At the 0.05 level, the population means are significantly different.

**Supplementary Table 22:** Descriptive Statistics for EndoGq Tracker's percent recovery in NCI-H125 vs HeLa cells without exogenous Gαq expression

| | N Analysis | Mean | Standard Deviation | SE of Mean |
|---|---|---|---|---|
| NCI-H125 | 9 | 86.05361 | 22.20409 | 7.40136 |
| HeLa | 10 | 74.84977 | 16.94655 | 5.35897 |

| | DF | Sum of Squares | Mean Square | F Value | Prob>F |
|---|---|---|---|---|---|
| Model | 1 | 594.59702 | 594.59702 | 1.54823 | 0.23028 |
| Error | 17 | 6528.84237 | 384.04955 | | |
| Total | 18 | 7123.43938 | | | |

At the 0.05 level, the population means are not significantly different.

**Supplementary Table 23:** Descriptive Statistics for recovery rates between EndoGq Tracker and PIP2 sensor in NCI-H125 cells

| | N Analysis | Mean | Standard Deviation | SE of Mean |
|---|---|---|---|---|
| EndoGq Tracker | 7 | 0.0031 | 0.00239 | 9.03697E-4 |
| PIP2 | 8 | 0.00716 | 0.00439 | 0.00155 |

| | DF | Sum of Squares | Mean Square | F Value | Prob>F |
|---|---|---|---|---|---|
| Model | 1 | 6.14295E-5 | 6.14295E-5 | 4.72253 | 0.04885 |
| Error | 13 | 1.69101E-4 | 1.30078E-5 | | |
| Total | 14 | 2.3053E-4 | | | |

At the 0.05 level, the population means are significantly different.

**Supplementary Table 24:** Descriptive Statistics for recovery extents between Gγ9 with and without EndoGq Tracker in NCI-H125 cells

| | N Analysis | Mean | Standard Deviation | SE of Mean |
|---|---|---|---|---|
| +EndoGq Tracker | 18 | 76.00178 | 7.66298 | 1.80618 |
| -EndoGq Tracker | 17 | 76.14933 | 8.31768 | 2.01733 |

| | DF | Sum of Squares | Mean Square | F Value | Prob>F |
|---|---|---|---|---|---|
| Model | 1 | 0.19036 | 0.19036 | 0.00298 | 0.95677 |
| Error | 33 | 2105.2033 | 63.79404 | | |
| Total | 34 | 2105.39366 | | | |

At the 0.05 level, the population means are not significantly different.

**Custom Macro Code used for Plasma membrane analysis in live cell imaging**

```
name = getTitle();
run("Duplicate...", "duplicate");
run("Enhance Contrast", "saturated=0.35 normalize stack");
run("Gaussian Blur...", "sigma=1.5 stack");
run("Find Edges", "stack");
setThreshold(x, y); // Set threshold to emphasize membranes depending on expression level
run("Convert to Mask");
run("Options...", "iterations=2 count=1 black");
run("Erode", "stack");
run("Dilate", "stack");
run("Analyze Particles...", "size=10-Infinity circularity=0.1-1.0 show=Nothing exclude clear add
composite stack");
selectImage(name);
run("Set Measurements...", "area mean stack redirect=None decimal=3");
roiManager("Measure");
```

**Legends for Supplementary movies**

**Supplementary Movie 1**

HeLa cell exhibits robust EndoGi Tracker recruitment to the plasma membrane upon $\alpha$2AR activation with norepinephrine.

**Supplementary Movie 2**

NCI-H125 cell exhibits robust EndoGq Tracker recruitment to the plasma membrane upon GRPR activation with bombesin. Interestingly, EndoGq Tracker recovers back to the cytosol despite continuous GRPR stimulation.

**Supplementary Movie 3**

HeLa cell expressing blue opsin alongside EndoGi Tracker exhibits localized EndoGi Tracker recruitment in response to localized 488 nm light activation. Blue box indicates light pulse. Upon exposing the whole cell to 488 nm light, EndoGi Tracker is recruited globally.
